# Tree shrews combine panoramic vision with an expanded oculomotor repertoire

**DOI:** 10.64898/2026.10.23.761344

**Authors:** Ratnadeep Pal, Erin Kempkes, Rishi Aeleti, Elizabeth Tubbs, Arnab Biswas, Madineh Sedigh-Sarvestani

## Abstract

Mammals differ in how they redirect gaze. Primates shift and hold gaze with rapid, precise eye movements that can be decoupled from the head. In lateral-eyed mammals studied to date, gaze is organized around the head: saccades are coupled to head rotation in freely moving animals and typically accompanied by attempted head movements even under head fixation. These observations suggest a trade-off: animals that keep a panoramic field give up the precise, eye-driven control of gaze typical of frontal-eyed animals. But because lateral eyes and head-centered gaze co-occur in every lateral-eyed species studied, it is unknown whether one follows from the other. Here we test this in the tree shrew, an arboreal mammal with laterally placed eyes. Using real-time eye tracking in head-fixed animals, we find that tree shrews saccade frequently under head fixation and that most saccades occur without detectable attempted head movement. Tree shrews make visually evoked saccades, along both the horizontal and vertical axes of the visual field, toward targets. They can also fixate on visual targets. Using widefield calcium imaging, we show that visual cortex over-represents the central visual field. Lastly, we show that extraocular muscles that move the eyes contain specialized nerve endings found in frontal-eyed species. Based on this suite of evidence, we conclude that lateral eyes do not preclude eye-driven gaze control. The two are separable dimensions of visual system design, a combination that may suit animals whose locomotion routinely points the head away from where they need to look.

---

Primates and other frontal-eyed animals explore the world with rapid, precise eye movements known as saccades. Saccades shift gaze to objects of interest, fixation holds it there, and both can occur independently of head movements^1,2^. In lateral-eyed mammals, gaze is more closely coordinated with movements of the head. One proposed explanation is that rotating a laterally placed eye adds little to an already wide monocular field that must be kept stable to maintain awareness of the surroundings^1,2^. In freely moving rodents, eye and head movements are strongly coupled. In mice, the eyes anticipate or compensate for head motion^3–5^, while in rats eye movements are likewise dictated by head movement and position and maintain broad binocular coverage rather than continuous fixation^6^. Classic accounts tie head-independent saccades to the evolution of foveae and frontal eyes^1,7^. Taken together, these observations suggest a broad trade-off: animals that keep a panoramic field give up the precise, eye-driven control of gaze that frontal-eyed animals enjoy.

Work in head-fixed mice adds nuance to this picture. Head-fixed mice make spontaneous and visually evoked saccades ^8–10^, but these movements are strongly coupled to attempted head movements^5,10^. This suggests that gaze remains anchored to the head even when the head is stationary. Head-restrained rats show a different signature of the same head-centered organization: their eyes move only infrequently (Fig. S1), even while they run on a treadmill^6^. The question, then, is not whether a lateral-eyed mammal can move its eyes, but whether it can move them as frontal-eyed mammals do: to shift gaze frequently and independently of the head, rapidly and toward visual targets, and then to hold gaze on a region of interest. Because nearly everything known about gaze in lateral-eyed mammals comes from rodents and rabbits, it remains unclear whether head-centered orienting follows from lateral eyes themselves or from the particular lineages studied. Furthermore, in primates, eye-driven gaze is one part of a coordinated phenotype that also includes a retina that concentrates sampling in a specialized region, a visual cortex that magnifies the central visual field, and extraocular muscles specialized for precise, voluntary control of eye position^11–13^. Whether a lateral-eyed mammal can assemble this combination of specializations while retaining a panoramic field is unknown.

Tree shrews offer a natural test of this question. They are diurnal and highly visual animals, evolutionarily close to primates, but classified into the order Scandentia. They are fairly small (average ∼150 grams) and have laterally placed eyes (Fig. 1A), with a binocular field of about 54 degrees within a total field exceeding 300 degrees^14,15^. They therefore pay the optical cost of lateral eyes through their relatively small binocular field. However, tree shrews are highly acrobatic animals who move rapidly through complex arboreal environments that demand fast, precise sampling of the visual scene. We therefore hypothesize that they may exhibit specializations for frontal vision, including eye movements decoupled from the head, despite their laterally placed eyes. Tree shrew eye movements have not been thoroughly characterized within their full range of natural motion. Recent measurements in freely moving tree shrews limited to planar locomotion during a chasing task revealed saccades tightly coordinated with head rotations^16^. However, this does not reveal whether the eyes can be controlled independently of the head during other behaviors that require the head and eyes to point in different directions. Measurement of eye movements during natural or artificial periods of head immobility can expose the capacity of head-decoupled gaze control.

**Figure 1:**
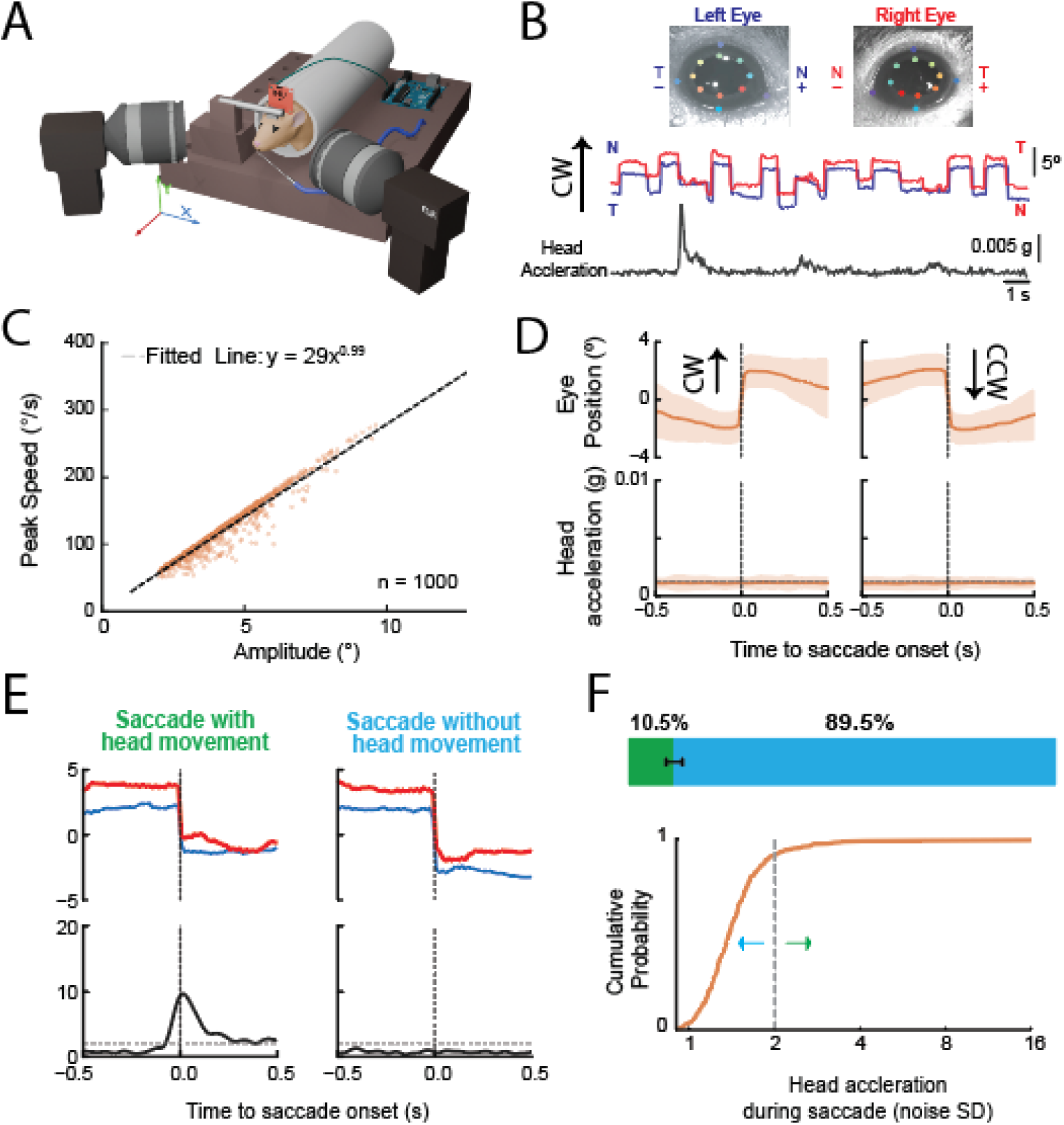
Tree shrews make head-independent saccades. **A)** Head-restrained eye and attempted head-movement tracking: Two infrared-sensitive cameras capture eye position, and an inertial measurement unit captures attempted head movements. **B)** Sample traces of horizontal eye position for the left and right eye, captured with DeepLabCut (top). Sample head acceleration magnitude traces (bottom). **C)** Saccadic main sequences for eye rotations about the yaw axis (n=1000 saccades in two tree shrews). **D)** Saccade-triggered average deviation of eye position for clockwise (CW) and counterclockwise (CCW) saccades (Top). Saccade-triggered average head acceleration magnitude (Bottom). The grey dashed line is the SD of the unfiltered accelerometer signal recorded with no animal (0.00127 g), shaded bands are SD. **E)** Representative traces of left and right eye position and head acceleration for saccades accompanied by attempted head movement (Left) and saccades without attempted head movement (Right). Head acceleration is in units of the SD of the unfiltered no-animal signal; the dashed line is the head movement threshold (2 units). **F)** Percentage distribution of saccade categories (Top), error bars are Wilson 95% CI. Cumulative probability of head acceleration magnitude within ±0.5 s of saccade onset (Bottom). n = 999 saccades from two tree shrews.

We developed a real-time eye-tracking setup for head-fixed animals that also records attempted head movements and combined these measurements with 3D models of the tree shrew eye in the head to determine how eye movements shift gaze through the visual field. We also used this setup, combined with a gaze-contingent display, to determine whether tree shrews can hold their gaze on a visual target. To determine the amount of cortical tissue devoted to central visual space, we used widefield calcium imaging. Finally, to investigate the biomechanical adaptations necessary for precise gaze control in the frontal visual field despite lateral orbits, we used histological methods to assess the muscles that move the eyes. We used rats as control for our methods for several analyses, as the oculomotor system of rodents is generally better established than tree shrews. We chose rats over mice because their closer size to tree shrews removes potential scaling confounds.

Together, we find that tree shrews combine lateral eyes with the oculomotor, cortical, and muscular specializations of frontal vision, revealing eye placement and gaze control as separable dimensions of visual system design.

## RESULTS

To determine whether tree shrews can make eye movements independent of head-movements, we developed a system for video based eye-tracking in head-fixed animals (Fig. 1A, see methods). The system included 2 high-resolution cameras, two infrared LEDs to improve imaging of the dark pupil, an inertial measurement unit (IMU) including a 3-axis accelerometer attached to the animal’s head to measure attempted head movements, and a juice delivery system. To validate that the head accelerations we measured with the head-mounted IMU reflected genuine attempted head movements, and that our IMU measurements did not miss small attempted head movements, we used an independent force-sensor apparatus^17^ that directly measured force generated at the head-fixation point along three axes (Fig. S1D). Across individual saccades, peak force-sensor signal was strongly correlated with peak IMU-measured head acceleration (Pearson’s r = 0.82; Fig. S1D), supporting the use of IMU measurements as an indicator of attempted head movement.

### Tree shrews make head-independent saccades

Using this system, we identified 70,520 horizontal saccades in two tree shrews passively viewing visual stimuli across 76 sessions and 22 hours and characterized their kinematics. Visual inspection of a random 1,098 saccades confirmed 1,091 (99.4%) of them as real saccades, as opposed to apparent eye movements due to poor eye tracking or camera motion. To maintain comparable numbers between tree shrews and rats (Fig. S1), who make far fewer saccades, the analyses below use a random n=1000 subset of the confirmed tree shrew saccades. Tree shrews made frequent eye movements while head-fixed, an average of 54.8 ± 12.3 saccades per minute (mean ± SD across sessions; median 53.7) during passive viewing. Saccade peak velocity scaled with amplitude according to a power-law relationship (Fig. 1C), consistent with the "main sequence" relationship that is a hallmark of saccadic eye movements across species. To determine whether these saccades occurred independently of attempted head movement, we analyzed head-acceleration near the time of eye movements using data from the IMU. Saccade-triggered averages showed a clear, stereotyped shift in eye position at saccade onset for both clockwise (CW) and counterclockwise (CCW) horizontal (yaw) saccades as expected. During these saccades, there was no corresponding increase in head acceleration above the noise floor (Fig. 1D), which was recorded with no animal on the rig. This indicates that, on average, tree shrew saccades are not accompanied by attempted head movement.

Averaging across all saccades could obscure a subset that co-occur with attempted head movements. To rule this out, we analyzed individual saccades and head-acceleration within ±0.5 s of each saccade and asked whether the animal attempted to move its head around the time of eye-movement (Fig. 1E). Saccades were categorized as “Saccades with head movement” if head acceleration crossed a threshold in this window, and as “Saccades without head movement” otherwise (Fig. 1E). We set the threshold as 2 times the standard deviation of the noise floor of the accelerometer. The threshold is deliberately low: a lower threshold labels more saccades as accompanied by head movement, which biases the analysis against our conclusion that saccades occur without it. We found that most tree shrew saccades occurred without a detectable head movement attempt (Fig. 1E, right). Across all n=1000 saccades, 89.5% (95% CI 87.4–91.2%) were categorized as a “Saccade without head movement” and only 10.5% (95% CI 8.8–12.6%) of saccades were categorized as a “Saccade with head movement” (Fig. 1F, top). Consistent with this, the cumulative distribution of head acceleration magnitude within ±0.5 s of saccade onset showed that many saccades were associated with head acceleration near the noise floor, well below the head-movement detection threshold (Fig. 1F, bottom). We also ran an automated analysis with every detected tree shrew saccade (n = 70,520). The results did not change: 9.6% of all detected saccades (95% CI 9.4–9.9%), against 10.5% of the confirmed saccades, occurred with attempted head movement. Saccades with attempted head movement were slightly larger than those without (all detections, median 4.2° vs 3.7°, p < 10⁻¹⁰⁰; confirmed saccades, median 4.1° vs 3.8°, p = 0.12), but the two distributions largely overlapped (Fig. S1E). Together, these results demonstrate that tree shrews can make saccadic eye movements independent of attempted head movement.

This near-complete independence of eye movements from attempted head movements in head-fixed tree shrews contrasts with head-fixed mice, in which eye movements are accompanied by attempted head movement^5,10^. To determine whether the uncoupling between the eyes and head that we observe in tree shrews is a general property of larger mammals, we performed the same eye-tracking and saccade-detection procedures in pigmented (Long-Evans) rats (Fig. S1A, n=2).

We found that head-fixed rats rarely move their eyes, producing a median of 0.51 saccades per minute, notably less than the median 53.7 in tree shrews measured under the same conditions. Rats also spent more time attempting to move their head (median 0.065 min per min, compared with 0.007 in tree shrews; Fig. S1A), and saccade-triggered averages in rats showed elevated head acceleration around the time of the saccade (Fig. S1B). Using the same head-movement threshold as in tree shrews, 61.7% (95% CI 56.4–66.8%) of individual rat saccades occurred with attempted head movement, compared with 10.5% in tree shrews (95% CI 8.8–12.6%; Fig. S1C, top). Nonetheless, the remaining 38.3% (95% CI 33.2–43.6%) of rat saccades occurred without attempted head movement, indicating that rats are capable of head-independent eye movements, at a rate that falls between the tight coupling reported in mice^4,5,10^ and the substantially higher rate we report here in tree shrews (89.5%; 95% CI 87.4–91.2%). Because these percentages depend on the choice of threshold, we also compared the cumulative distribution of head acceleration during saccades in the two species (Fig. S1C, bottom). The rat distribution is shifted toward higher head acceleration across its entire range, so regardless of where the threshold is placed, a smaller fraction of rat saccades than tree shrew saccades would be classified as head-independent. These results indicate, perhaps unsurprisingly, that the head-driven gaze strategy predominant in rodents cannot be explained merely by body or head size, nor by the shared lateral position of the eyes in the orbit. Instead, they point to a genuine species-specific difference in oculomotor control, with tree shrews exhibiting a more complex gaze-control strategy consistent with their complex movement repertoire and visuomotor needs.

### Tree shrew saccades produce horizontal and vertical gaze-shifts in the frontal field

Having established that tree shrews can make saccadic eye-movements independent of head-movements, we next asked about the consequence of these eye movements on sampling of the visual field. Specifically, we set out to study the mapping between the axis of rotation of the eye and the axis of gaze-shift in the visual field. This mapping requires accounting for the tree shrew’s lateral orbits (Fig. 2A), which unlike frontal-eyed primates (Fig. 2B), creates a divergence between the eye’s optical and visual axis. The optical axis is the line through the center of the pupil, whereas the visual axis is the line of sight defined by the projection of the retinal specialization into the visual field. The retinal specialization in tree shrews lies deep into the temporal retina^18^ (Fig. 2A), directing the visual axis forward, into the binocular visual field.

**Figure 2:**
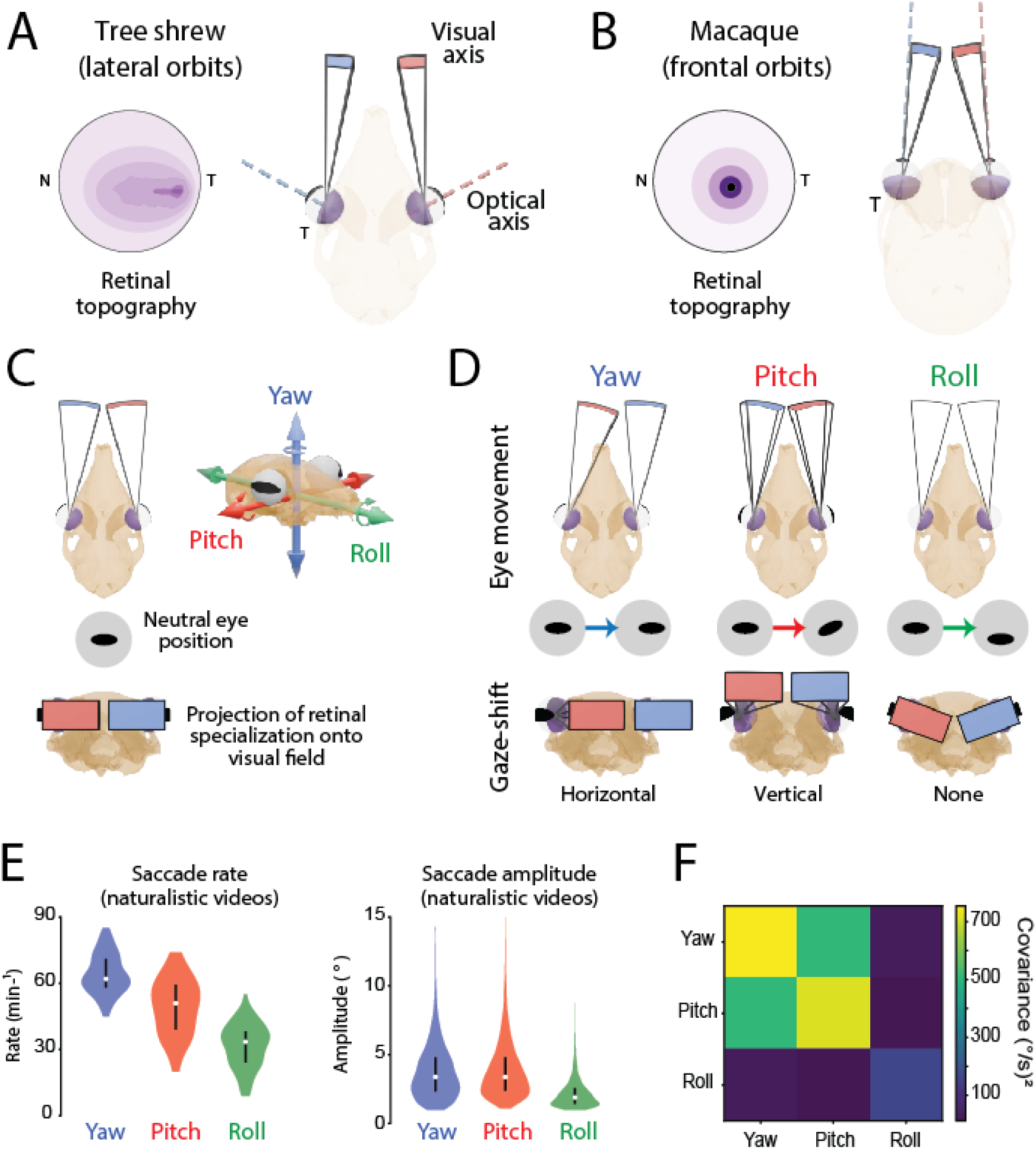
Tree shrew saccades produce horizontal and vertical gaze-shifts in the frontal visual field. A) Tree shrews have laterally placed eyes/orbits and a retinal specialization comprising a temporal area centralis and a horizontal streak^18^. Retinal shading schematically represents relative ganglion cell density (approximately 3,000–20,000 cells/mm²). The lateral placement of the eyes produces a divergence between the optical axis, passing through the center of the eye, and the visual axis, projecting from the retinal specialization. This produces a divergence in their optical axis: the line through the center of the pupil and their visual axis: the line projecting from the retinal specialization. **B)** Macaques and many primates have frontal eyes/orbits, and a central retinal specialization (fovea), which produces approximately aligned visual and optical axes. Retinal shading schematically represents relative ganglion cell density (approximately tens to 50,000 cells/mm²)^19^. Shading in A and B is normalized independently for each species, with darker colors indicating higher relative density. **C)** Top-view of tree shrew skull + eye showing neutral eye position and side-view showing three axes of eye movements. Approximate projection of retinal specialization onto the visual field is shown in red and blue for the left and right eye, respectively. **D)** The three types of eye movements, shown in top-view (top), from the perspective of a camera parallel to the eye (middle), and front-view showing resulting impact on gaze-shift in visual field (bottom). **E)** Saccade rate (left) and amplitude (right) for the three types of eye-movements, recorded while animals (n=2) passively viewed naturalistic movies. Data are pooled across 5 sessions and include 5,301 yaw, 4,059 pitch and 2,627 roll saccades. **F)** Covariance of eye velocity among the three axes (deg/s)², over all frames of both eyes, pooled across both animals.

The divergence between the optical and visual axis in tree shrews and other lateral-eyed animals matters for how eye movements are measured and interpreted. Eye-tracking cameras are typically placed parallel to the eye, and therefore capture pupil displacement along the optical axis, not the visual axis. When these axes diverge significantly as they do in tree shrews, tracking the pupil does not equate to tracking gaze. One solution is to place eye-tracking cameras parallel to the nose, and perpendicular to the visual axis. But it would be difficult to capture the whole eye from this vantage point. We therefore placed eye-tracking cameras in the standard configuration, approximately parallel to the eyes (Fig. 1A), and built geometric models of the eyes within their skeletal orbits to predict how rotations about different axes appear in camera space and shift gaze in the visual field (Fig. 2; Fig. S2; Methods). To formalize this, we considered the three rotational axes of eye movement: yaw, pitch, and roll (Fig. 2C), in a head-based reference frame . For each axis, we simulated the consequence of eye movement on a) the eye-tracking camera placed parallel to the eye, and b) the gaze-shift produced in the visual field. To help build our intuition, we also built a 3D model of macaque eyes in their skeletal orbits (Fig. 2C-D, Fig. S2A-D).

Yaw rotations are orthogonal to the optical axis in both frontal & lateral configurations and accordingly appear as horizontal pupil displacement on camera and produce horizontal gaze-shifts in both lateral and frontal-eyed species (Fig. 2D, Fig. S2C-D). Along pitch and roll, the impact of eye rotations on gaze diverges for frontal vs. lateral-eyed species. Because the tree shrew orbit points laterally, pitch rotations produce pupil torsion, an angular rotation of the eye along the optical axis, which produces rotation but no translational displacement of the pupil on camera (Fig. 2D, S2A). Such torsion is difficult to detect in round-pupiled animals like mice but is readily measured in elliptical-pupiled tree shrews via the changing orientation of the pupil’s long axis. Despite this lack of translational displacement, pitch rotations produce a vertical gaze-shift in tree shrews. Roll rotations on the other hand, produce vertical pupil displacement on camera but little gaze-shift, instead simply rotating the retinal image in place (Fig. 2D, S2D). In macaques, this pattern is reversed: the optical axis aligns with the naso-occipital axis, so it is roll rather than pitch that produces torsion on camera, and because the visual axis is centered on rather than offset from the optical axis, this rolling motion produces only rotation of the retinal image (Fig. 2D, S2B).

This modeling exercise yields an anatomically grounded prediction: Tree shrews should preferentially recruit yaw and pitch eye movements during natural visual behavior and largely avoid roll, since only yaw and pitch produce gaze-shifts. In other words, we should see clear horizontal displacement of the pupil (yaw), and clear rotation of the pupil along the long-axis of the ellipse (pitch), but relatively little vertical displacement of the pupil (roll). We tested this prediction in head-fixed tree shrews (n=2) freely viewing naturalistic videos, classifying saccades by their dominant rotational axis (see Methods). Consistent with our prediction, tree shrews made yaw and pitch saccades at substantially higher rates than roll saccades (Fig. 2E, left; Fig. S2E), and yaw and pitch saccades were also of larger amplitude than roll saccades (Fig. 2E, right; Fig. S2F). The median saccade rates were 62, 51 and 33.5 saccades per minute for yaw, pitch and roll, and the median amplitudes were 3.4°, 3.4° and 1.9°, respectively. In addition, yaw and pitch eye velocity varied together, whereas roll velocity was nearly independent of both (covariance over all frames: yaw–pitch 505 (deg/s)², yaw–roll 29, pitch–roll 12; Fig. 2F) Together, we find that the tree shrew’s saccadic repertoire is structured around the two eye-movement axes effective at shifting gaze horizontally and vertically in the visual field.

### Head-fixed tree shrews make visually evoked directional saccades

We have established that tree shrews make head-independent saccades, along axes that produce horizontal and vertical gaze-shifts in the frontal binocular field. However, spontaneous saccades made while viewing naturalistic scenes cannot distinguish random or reflexive eye movements from the voluntary, target-driven oculomotor control that characterizes frontal-eyed species. If tree shrews use saccades to place their retinal specialization on objects of interest, then saccades should be evoked by visual stimuli and directed toward them. We tested this with a saccade-to-target task.

Tree shrews orient their eyes toward newly presented visual targets without training, but these responses are inconsistent and easily interrupted by competing stimuli. We therefore trained animals through progressively more demanding reward contingencies to produce target-directed saccades reliably across many trials (see Methods). In the final task, each trial began with a central cue followed by a target appearing randomly on the left or right side of a monitor placed in the frontal visual field (Fig. 3A). Trials were considered valid if a saccade occurred within 1 second after target onset. Valid trials were considered correct and rewarded if the first saccade was in the direction of the target, with a tolerance of ±90°.

**Figure 3:**
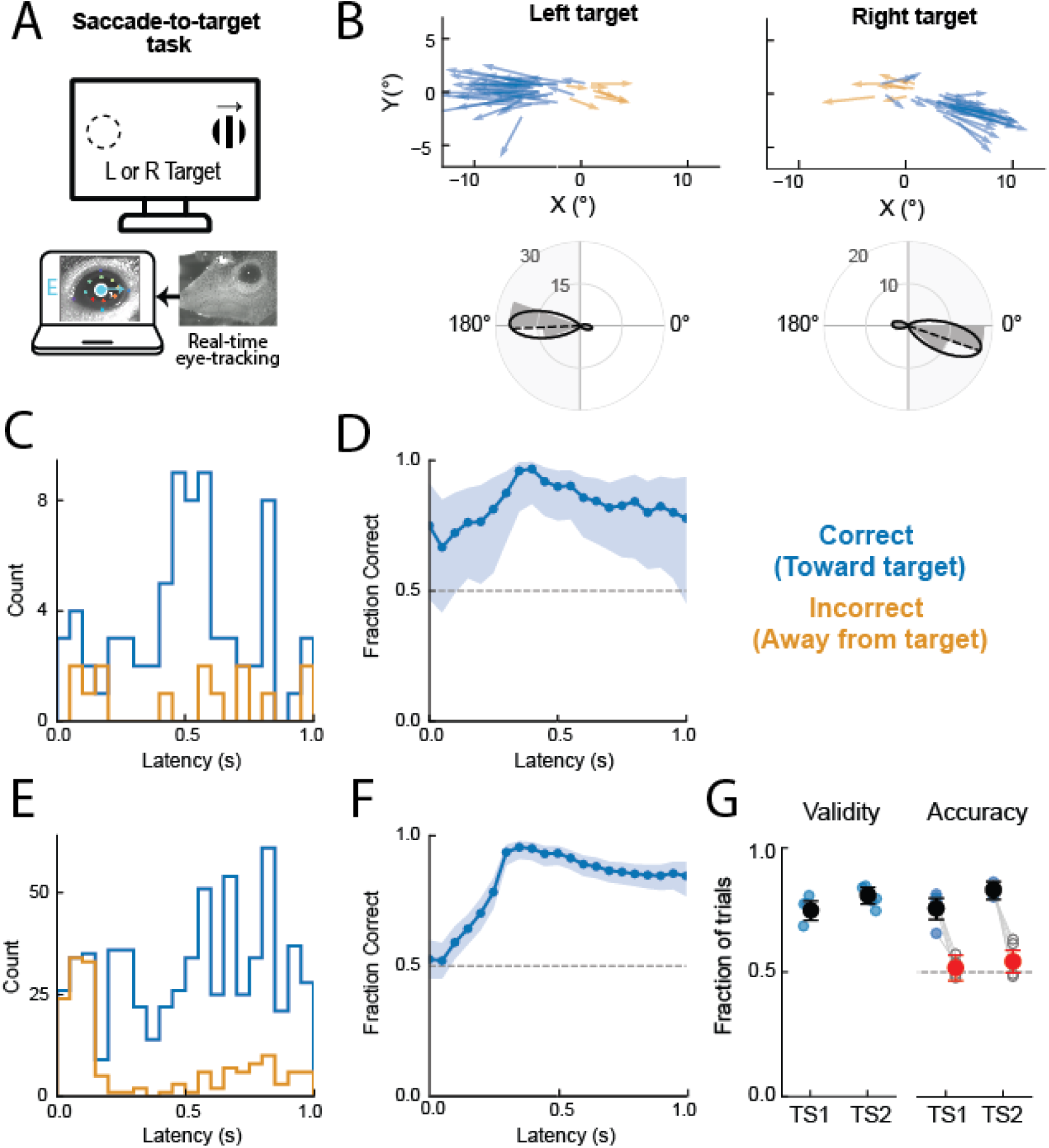
Head-fixed tree shrews make visually evoked directional saccades. **A)** Saccade-to-target task. A central visual cue is followed by a target appearing randomly on the left or right side of the monitor. Trials are classified as valid if the animal makes a saccade within 1 s of target onset, and invalid otherwise. Valid trials are scored as correct if the *first* saccade after target onset is directed toward the target and incorrect if directed away from it. Correct trials are rewarded. **B)** First-saccade trajectories in monitor coordinates (top) and corresponding polar distributions of saccade direction (bottom) for left-target (n = 48 valid trials) and right-target (n = 39 valid trials) conditions in an example session. Blue arrows show correct and orange arrows incorrect saccades. In polar plots, bars show number of saccades in each 20° direction bin , the black curve is a smoothed distribution, the dashed line marks the circular mean direction, and shaded sectors mark the directions scored correct. **C)** Distribution of first-saccade latencies relative to target onset in the example session (n = 87 valid trials). **D)** Fraction of correct first saccades in 0.3 s sliding windows for the same session (n = 87 valid trials). Blue shading indicates pointwise 95% Wilson confidence intervals; dashed line indicates chance (0.5). **E–F)** Same as C–D, pooled across 10 sessions from two animals (five sessions per animal; n = 801 valid trials). **G)** Performance across all sessions. Validity (left): fraction of all trials containing a saccade within 1 s of target onset. Accuracy (right): fraction of correct saccades compared with a pre-cue control (open symbols). Individual points are sessions; black (validity and accuracy) and red (pre-cue control) markers are trial-pooled estimates for each animal with 95% Wilson CIs.

In an example session, saccades were strongly biased toward the target. Left targets predominantly evoked leftward saccades, and right targets predominantly evoked rightward saccades (Fig. 3B). To determine whether these saccades were visually evoked, we examined their onset time relative to target onset. Correct saccades became more frequent following target onset (Fig. 3C), and the fraction of correct saccades increased sharply beginning around 0.2 s, reaching approximately 90% at latencies of 0.3–0.4 s (Fig. 3D). This pattern was preserved across both animals (five session per animal; Fig. 3E–F, Fig. S3B). At the shortest latencies (0–0.2 s), saccade direction was close to chance, suggesting that animals were guessing prematurely prior to accumulating sufficient visual evidence. The latencies we report here are longer than those typically reported for trained primates^20^, although our task did not require central fixation before target onset, and differences in starting eye position, attentional state, and training may contribute to the longer responses.

Although most tree shrew saccades occur without attempted head movement (Fig. 1F), saccades made during a task, and correct saccades in particular, might still recruit the head. To test this possibility, we classified first saccades from a subset of sessions recorded with the head-mounted IMU, using the same criterion as in Fig. 1. The fractions of saccades with and without head movement were similar to those during passive viewing (9.4% vs 10.5% with head movement; Fig. 1F), and correct saccades were no more likely to involve head movement than incorrect saccades (8.2% for correct vs 11.9% for incorrect; Fig. S3A). This suggests that, similar to saccades made during passive viewing of visual stimuli, saccades made in the saccade-to-target task occur largely independent of an attempted head movement.

We quantified performance across sessions using two measures: validity, the fraction of all trials containing a saccade within 1 s of target onset, and accuracy, the fraction of correct saccades. Across five sessions per animal, 77.0% of trials were valid in TS1 (379/492) and 80.7% in TS2 (422/523). Of these valid trials, 75.7% were correct in TS1 (287/379) and 83.2% in TS2 (351/422; Fig. 3G). To determine if saccade direction depended on target location, we compared accuracy with a pre-cue control, in which saccades occurring before target onset were scored relative to the upcoming target direction. Pre-cue saccades were approximately equally likely to be directed toward or away from the upcoming target (51.8% in TS1 and 54.4% in TS2), in contrast to the strong directional bias following target onset (Fig. 3G). These results demonstrate that visual target appearance influences both the timing and direction of saccades in head-fixed tree shrews.

Finally, we asked whether vertically displaced targets could evoke the corresponding pitch saccades, which our eye-in-head model predicts should appear on camera as torsion of the elliptical pupil rather than translational pupil displacement (Fig. 2D). Indeed, upward targets reliably evoked clockwise (CW) torsion saccades—the camera signature of an upward pitch rotation—at short latencies after target onset (Fig. S3C–E). Downward targets, however, did not evoke the complementary torsion saccades, possibly because the tree shrew’s long snout occludes the lower frontal field where the downward target appeared. Together, these results demonstrate that head-fixed tree shrews can direct visually evoked saccades along both axes of the frontal binocular field, providing a means to aim their temporal retinal specialization toward visual targets despite their laterally placed eyes.

### Head-fixed tree shrews can fixate a visual target

Saccades solve only half the problem of frontal visual specialization: relocating a small high-acuity region of retina onto a target is of little use unless gaze can then be stabilized on that target long enough to extract detailed information. This ability, referred to as fixation, is a defining feature of frontal-eyed oculomotor control necessary for precise visual inspection. Having shown that head-fixed tree shrews can direct their gaze toward a visual target, we next asked whether they can hold it there — that is, whether tree shrews can fixate. We tested this in two stages: first asking whether tree shrews can voluntarily prolong periods of stable eye position on cue (Fig. 4A–C), and then whether they can hold gaze at a specific location in the visual field (Fig. 4D–G).

**Figure 4:**
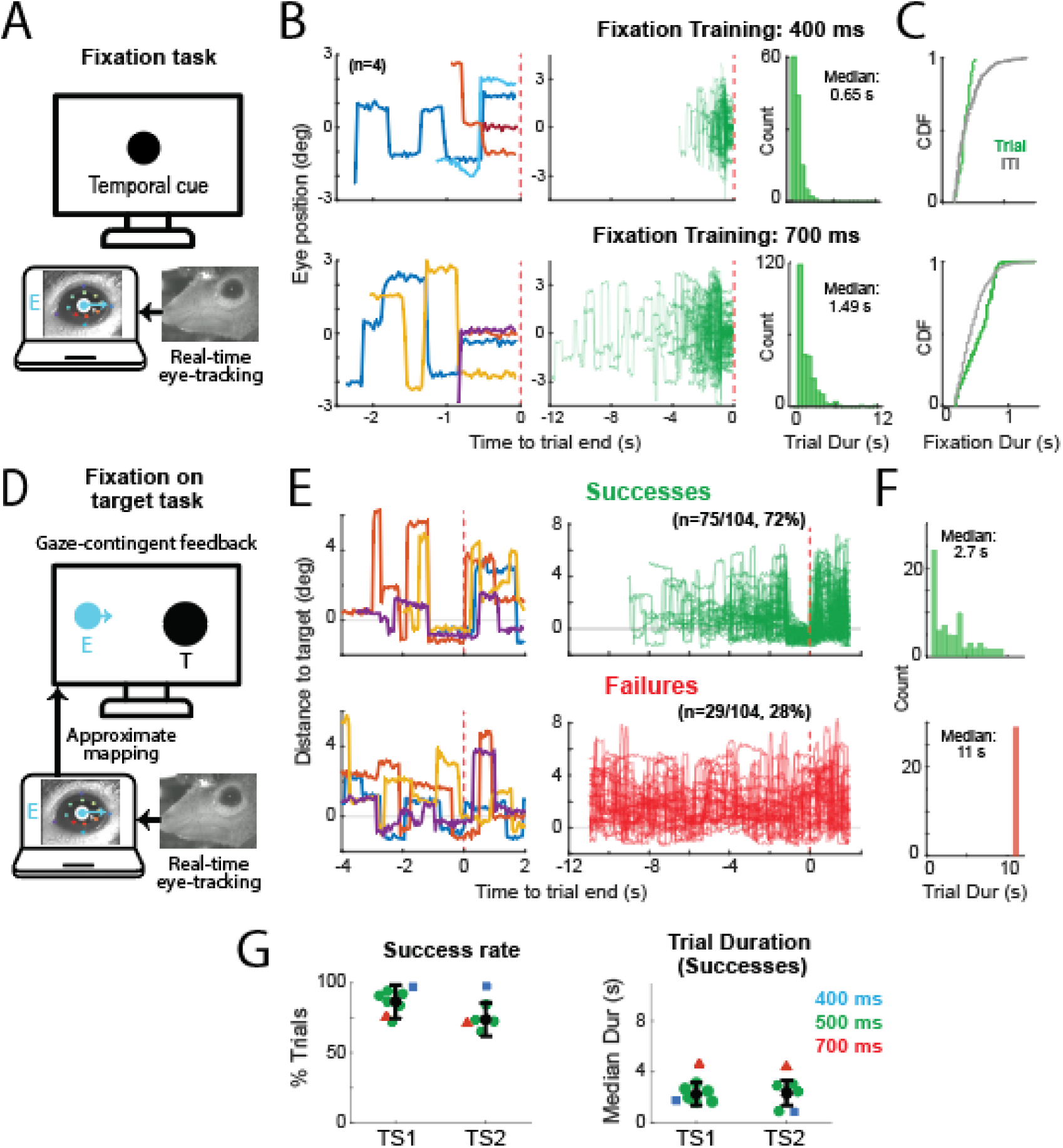
Head-fixed tree shrews can fixate. **A)** Fixation task used for cued fixation-training. A visual cue appears on the monitor and the animal is rewarded for fixations of increasing duration (300-750 ms), which end the trial. **B)** (top row) Horizontal eye-position for n=4 sample trials of a 400 ms fixation session, zoomed in on the last 2.5 seconds before trial- end (left), for 100 randomly chosen trials (of 127) and for the full trial duration (middle), and histogram of all trial durations (right). Bottom row shows the same data for a session with 700 ms fixation. **C)** Cumulative distribution function (CDF) of fixation duration for all trials (green), and for fixations calculated from the inter-trial-interval (gray) for the 400 ms session (top) and 700 ms session (bottom). **D)** Fixation-on-target task used for training and testing of on-target fixation. The animal’s gaze on the monitor is estimated and presented using a blue marker (E) which refreshes in real time. A black visual ‘target’ (T) appears in randomized order in one of two positions. The animal is rewarded, for bringing the blue gaze-contingent marker to the black target cue and holding fixation for 500 ms. If on-target fixation has not been achieved in 11 seconds, the trial fails without reward. **E)** (top row) Distance to target, in degrees, for n=4 sample successful trials in a sample session, zoomed in to 6 seconds around trial-end (left) and for all successful trials (n= 75). Bottom row shows the same data for failed trials. **F)** Histogram of trial duration for successful (top) and failed (bottom) trials. **G)** Population results for two animals. Each marker indicates a session. Markers are colored by fixation duration.

We first trained animals on a cued fixation task (Fig. 4A). On each trial, a visual cue appeared on the monitor, and the animal was rewarded for holding its eye position stable — without saccades — for a criterion duration between 300 and 750 ms, which increased with training; completion of the criterion fixation ended the trial. Trial duration therefore measures how quickly the animal produced the required fixation after cue onset. In example sessions with 400 ms and 700 ms criteria, horizontal eye position was held stable in the final portion of each trial (Fig. 4B), and trials were short, with median durations of 650 ms and 1490 ms that scaled with the criterion (Fig. 4B, right) — animals produced criterion fixations promptly after the cue rather than accumulating them by chance as time passed. Furthermore, to determine whether animals could control the duration of fixations, we compared the distribution of fixation durations during trials against fixation durations computed from the inter-trial interval, when no fixation was required (Fig. 4C). At the 400 ms criterion, the two distributions largely overlapped, indicating that fixations of this duration fall within the range produced by the animals’ spontaneous oculomotor behavior. At the 700 ms criterion, however, within-trial fixation durations were clearly shifted toward longer values than inter-trial fixations (median 0.57 against 0.43 s; Mann-Whitney U test, p<<0.05). This suggests that when the task demanded fixations longer than those occurring spontaneously, tree shrews extended them accordingly. This adjustment of fixation duration to task demands, together with the prompt timing of criterion fixations after cue onset, suggests that tree shrew fixation is under voluntary control rather than a passive byproduct of infrequent saccades.

Cued fixation demonstrates control over *when* the eyes are held still, but not *where*. Testing whether tree shrews can fixate a visual target requires knowing precisely where the animal is looking on the stimulus monitor, but the calibration needed to map eye position onto the monitor with such precision is too lengthy to perform each session. We therefore closed the loop through the animal instead. Gaze position on the monitor was estimated in real time from an approximate eye-to-monitor mapping and displayed to the animal as a blue marker (Fig. 4D). Because the reward contingency was defined on this visible marker rather than on the animal’s true (unknown) gaze position, imprecision in the mapping did not compromise the task: the animal simply had to learn to steer the marker onto the target, whatever the mapping between its eye movements and the marker happened to be. On each trial, a black target appeared at one of two positions in randomized order, and the trial was scored a success, and rewarded, if the animal brought the gaze-contingent marker to the target and held it there for 500 ms; trials without on-target fixation ended in failure after 11 seconds.

In an example session (Fig. 4E), 75 of 104 trials (72%) were successful. On successful trials, the distance between estimated gaze and target converged toward zero in the final moments before trial end, whereas on failed trials gaze wandered without settling on the target. Importantly, on-target fixations did not arise from chance encounters between a wandering gaze and the target: heatmaps of gaze position on the monitor showed that gaze density concentrated around the target and tracked its location as target position varied— a pattern present across all trials, including failures (Fig. S4). Even when animals failed to achieve criterion fixation, they allocated gaze around the target location, indicating that gaze was directed at the target rather than coinciding with it accidentally. Successful trials were completed promptly (median duration 2.7 s), well before the 11-second timeout (Fig. 4F). Furthermore, we note that success at this task requires not only fixation but learning to use visual feedback about one’s own eye position — a hallmark of flexible, voluntary oculomotor control.

Across 14 sessions in both animals, success rates were maintained as the required fixation duration increased from 400 to 700 ms, with median trial durations for successful trials remaining short across criteria (Fig. 4G). Together, these results show that tree shrews possess key features of the oculomotor repertoire of frontal vision: they rapidly shift gaze to a target with a visually evoked saccade (Fig. 3) and then voluntarily hold it there (Fig. 4), demonstrating a combination of abilities previously associated with frontal-eyed mammals.

### Tree shrew visual cortex over-represents the central visual field

Shifting and holding gaze with the eyes is useful only if some part of the retina is worth aiming at relevant portions of the visual field. In primates, the fovea is a highly specialized region of the retina with a high density of photoreceptors, whose representation is greatly magnified in primary visual cortex (V1). Tree shrews lack a fovea, but possess a region of elevated cell density in the temporal retina that projects to the frontal visual field(Fig. 2A)^18^. If tree shrews use saccades and fixation to direct this retinal specialization toward objects of interest, we might expect their visual cortex to devote a disproportionate share of its surface area to the central visual field. We tested this by measuring the cortical representation of the central 10° in tree shrew V1.

We imaged GCaMP6s responses through a large cranial window, across the surface of V1 in awake animals viewing retinotopic-mapping stimuli (Fig. 5A). A bar swept across the frontal visual field along the horizontal and vertical axes, and the phase of the evoked response at each cortical location was used to construct maps of azimuth and elevation^24^ (Fig. 5B). These maps revealed orderly representations of visual space in both hemispheres, with the vertical meridian running obliquely across V1 and lower elevations represented more anteriorly, consistent with previous electrophysiological and imaging studies^25–27^. After removing vascular artifacts, we used these maps to identify the portion of V1 representing the central 10° of the contralateral visual field, defined as 0–10° azimuth and −10° to 10° elevation (Fig. 5B,D).

**Figure 5:**
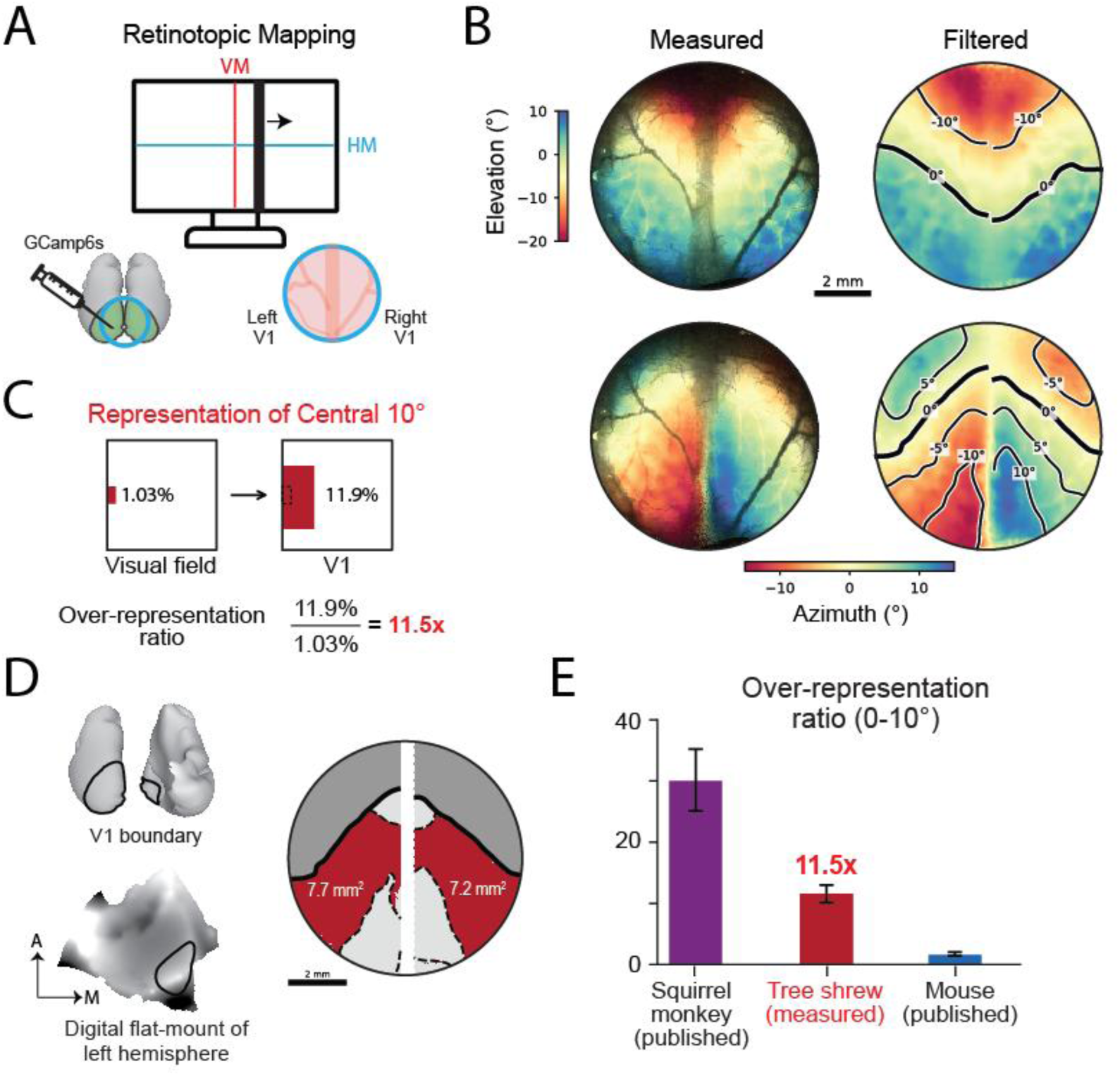
Tree shrew visual cortex over-represents the central visual field (0-10°). **A)** Retinotopic maps were measured in V1 using widefield GCaMP6s calcium imaging in awake animals passively viewing stimuli. A bar drifted across the monitor, rightward to map azimuth and upward to map elevation (VM, vertical meridian; HM, horizontal meridian). **B)** Maps of elevation (top) and azimuth (bottom) in the V1 imaging window: left, as measured; right, after removal of vessel artefacts, with contour lines. **C)** Schematic to show over-representation ratio of the central 10°, which compares fraction in the visual field to fraction in V1 surface. Squares represent total visual field coverage (left) and total V1 surface area (right). The red rectangle in each represents the central 10° has (1.03% of the visual field, 11.9% of V1). The dashed black line is a depiction of V1 surface area for uniform representation (ratio =1). **D**) (Left) We used digital flattening of a 3D tree shrew brain, published as part of a digital anatomical atlas^21^, to calculate the total surface area of V1 (Methods). (Right) Portions of V1 within our imaging window that fell within the central 10 degrees (0-10 azimuth, -10 to 10 elevation). The red region reaches the edge of the window, which holds 69% and 80% of the contralateral half of the central 10° in the two hemispheres. **E)** Over-representation ratio of the central 10° for three species. The ratio for squirrel monkey^22^ , > 30, and mouse, 1.6, are computed from published data^23^. Error bars represent values obtained under alternative definitions, which differ by species (Methods).

To quantify the degree of central-field magnification, we calculated an over-representation ratio: the fraction of V1 surface area devoted to a region of visual space divided by the fraction of the total visual field occupied by that region (Fig. 5C). A ratio of 1 indicates uniform allocation of cortical surface across the visual field, whereas values greater than 1 indicate disproportionate representation. We first estimated the total surface area of V1 using a digital flat-mount reconstructed from a published atlas of the tree shrew brain^21^ (Methods, Fig. 5D). This yielded a V1 surface area of 62.5 mm². Within our imaging window, the cortical representation of the central 10° occupied 7.7 and 7.2 mm² in each hemispheres, or 11.9% of total V1 surface area on average.

Our imaging window did not capture the entire representation of the central 10°, covering approximately 69% and 80% of this region in the two hemispheres (Fig. 5D). We therefore restricted our calculation to the portion of visual space represented within the window. Using published electrophysiological measurements to estimate the full extent of the contralateral visual field represented in V1²⁴ (Fig. S5A), we found that the exposed portion of the central 10° occupied only 1.03% of the total visual field, despite accounting for 11.9% of V1. This corresponds to an over-representation ratio of 11.5, with ratios of 12.8 and 10.4 in the two hemispheres (Fig. 5C,E). Thus, the central visual field receives approximately eleven times more cortical surface area than expected from a uniform representation of visual space. Because the precise location of the horizontal meridian depends on the alignment of the eyes relative to the stimulus monitor, we repeated the calculation after shifting its assumed position by ±5°. The resulting ratios ranged from 10.1 to 13.0, indicating that substantial central-field over-representation is robust to this uncertainty (Fig. 5E).

To independently assess this estimate, we repeated the analysis using previously published measurements of tree shrew V1^25,26^ (Fig. S5). From the retinotopic map of Bosking et al.^25^, we estimated that the full central 10° occupies approximately 9.5 mm² of cortex (Fig. S5B). After accounting for the fraction of this region visible through our cranial window, the predicted cortical areas closely matched our measurements in both hemispheres (Fig. S5C). Combining this published map with an independent estimate of total V1 surface area of 66 mm^28^ and the visual-field coverage reported by Kaas et al.^26^ yielded an over-representation ratio of 10.3 (Fig. S5D), in close agreement with our directly measured value of 11.5.

To place the tree shrew results in context, we compared the degree of central 10° over-representation with estimates we calculated from published retinotopic maps of squirrel monkey^22^ and mouse^23^ (Fig. 5E). We chose squirrel monkeys because they are a foveated primate for which exceptionally precise measurements of V1 retinotopy are available^22^. The central visual field was over-represented by approximately 30-fold in squirrel monkeys (25–35 across alternative estimates of the binocular field boundary), compared with 11.5-fold in tree shrews and only 1.6-fold in mice (1.4–2.0 across alternative definitions of the visual-field center). The mouse estimate was derived from published imaging data^23^, which may yield reduced estimates of visual field coverage, inflating the over-representation ratio. An independent electrophysiological map^29^ yielded an even lower ratio of 0.8 (range 0.5–2.2). The whiskers in Fig. 5E reflect uncertainty in the anatomical or visual-field reference positions used for these calculations, rather than variability between animals (see Methods). Thus, tree shrew V1 over-represents the central visual field approximately seven times more strongly than mouse V1, and notably less than the foveate squirrel monkey.

Together, these results show that tree shrews devote a disproportionate share of visual cortex to the frontal visual field, despite their laterally placed eyes and panoramic visual coverage. This cortical specialization complements the oculomotor abilities demonstrated in Figures 1–4: tree shrews can not only shift and hold gaze on visual targets but also allocate substantial cortical resources to the region of visual space toward which those movements are directed.

### Tree shrew extraocular muscles have morphologically elaborate sensorimotor innervations typical of frontal-eyed animals

Precise control of eye movements depends on the extraocular muscles (EOMs), which are innervated by the axons of neurons in the brainstem oculomotor nuclei (Fig. 6A). Conventional motor terminals form neuromuscular junctions along the muscle fibers, with large motor endplates concentrated near the middle of the muscle and smaller terminals distributed along certain fibers. In addition to these motor terminals, some EOMs contain specialized nerve endings called palisade endings, located at the distal myotendinous junction where the muscle meets the tendon (Fig. 6B). Here, a nerve fiber reaches near the end of the muscle fiber, sometimes turns back, and forms a cuff of terminal branches around the tip of a muscle fiber (Fig. 6B). Although palisade endings have features associated with motor innervation, their function remains unresolved^30–33^.

**Figure 6:**
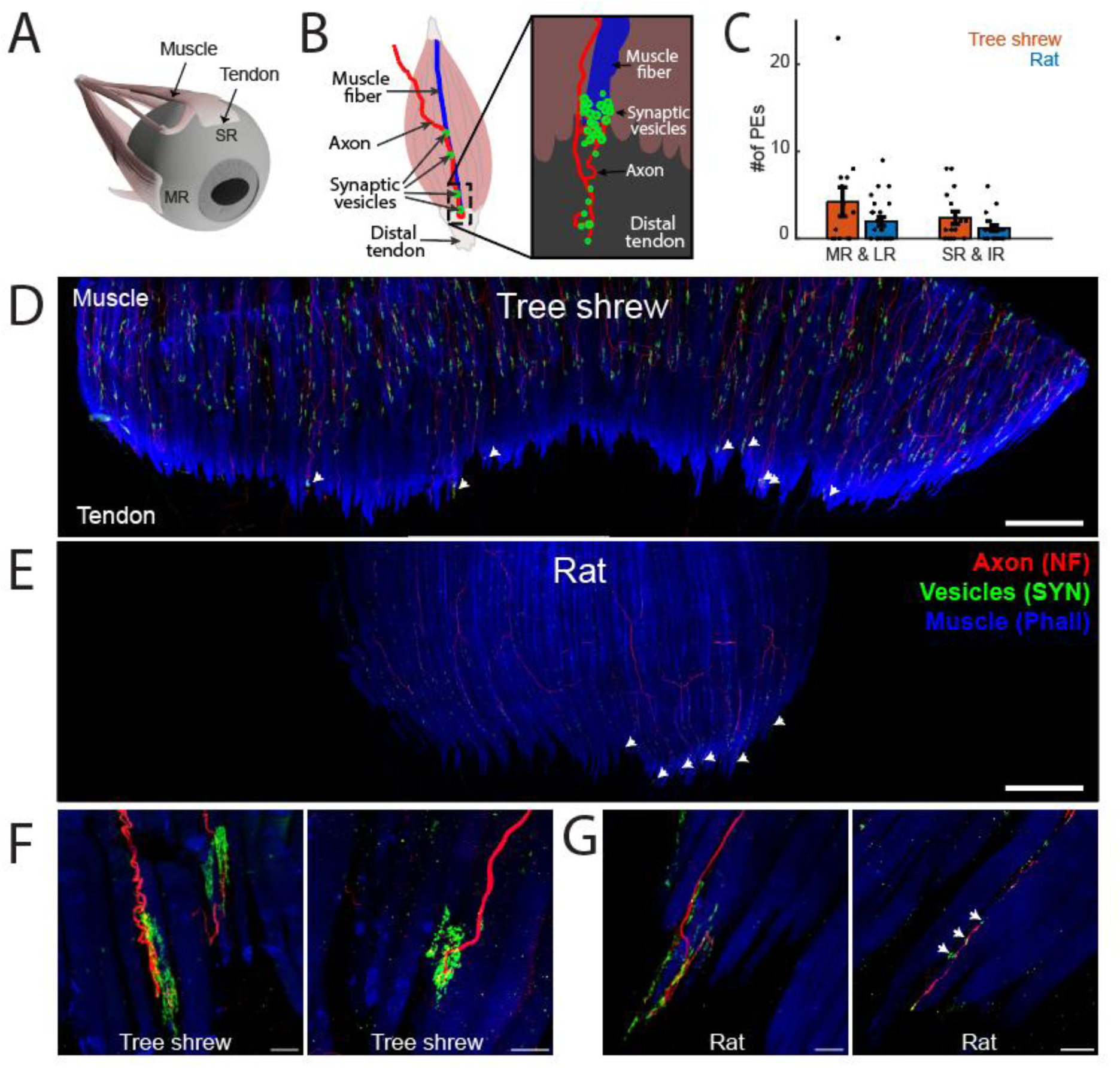
Tree shrew extraocular muscles have more numerous and elaborate sensorimotor innervations than rat extraocular muscles. **A)** Schematic of the eye and extraocular muscles, showing muscle and tendon. The horizontal recti (medial (MR) & lateral (LR)) contribute to horizontal gaze shifts whereas the vertical recti (superior (SR) & inferior (IR)) produce vertical gaze shifts. **B)** Schematic of an extraocular muscle. Box shows a palisade ending (PE), a specialized sensorimotor innervation, located at the distal myotendinous junction. A nerve fiber (red) forms terminal branches around the tip of a muscle fiber (blue). Adapted from^33^. **C)** Number of PEs per muscle in the horizontal/vertical recti of tree shrews (14 horizontal and 16 vertical from 4 animals) and rats (25 horizontal and 21 vertical from 7 animals). Bars indicate mean ± SEM. **D-E)** Z-projection of confocal tile scan of the entire distal myotendinous junction of a tree shrew (D) and rat (E) LR muscle labeled for neurofilament (NF, red), synaptophysin (SYP, green) and phalloidin (Phall, blue). Palisade endings (PEs) are noted with arrows. Scale bar, 500 µm. **F-G)** Higher magnification confocal z-projected image of example palisade endings in a tree shrew (F) and rat (G) EOM. Scale bars, 25 µm. In both panels, the image on the left shows an example of a highly developed PE for that species, while the image on the right shows an example PE with more simple morphology. Arrows in G) (right) indicate synaptophysin-positive terminals alongside the muscle fiber.

In a survey of 13 mammalian species, palisade endings were consistently found in the rectus muscles of all frontal-eyed species examined, including primates and carnivores^13^. In lateral-eyed species, they were generally less numerous or absent, and their morphology was simplified. Notably, rats exhibited relatively few palisade endings with simplified terminal branching, whereas mice lacked them entirely^13^. Given the relatively limited oculomotor repertoire of rodents, this pattern suggests that palisade-ending organization may be related to oculomotor capacity. Supporting this possibility, experiments in developing cats showed that eye immobilization reduced both the number and morphological complexity of palisade endings^34^. Here we asked whether tree shrew EOMs contain palisade endings resembling those of frontal-eyed mammals or the simpler form described in rats. We chose rats for comparison because we had also characterized their eye movements under head fixation (Fig. S1), and because their previously documented palisade endings provided a positive control for our labeling and imaging methods.

We dissected the four rectus muscles (Fig. 6A) from tree shrews (n = 4) and rats (n = 7), labeled them for neurofilament (NF) to mark axons, synaptophysin (SYP) to mark nerve terminals, and phalloidin to mark muscle fibers, and imaged the myotendinous junction with confocal microscopy (see Methods). Palisade endings were identified as NF- and SYP-positive terminal arborizations at the tips of muscle fibers. We found palisade endings at the myotendinous junction of EOMs in both species (Fig. 6D–G). In rats they were sparse and absent from many muscles, averaging 1.9 ± 0.5 per horizontal rectus and 1.2 ± 0.3 per vertical rectus (mean ± SEM; n = 25 horizontal and 21 vertical muscles), within the range previously reported^13,35^. In tree shrews they were more numerous, averaging 4.2 ± 1.7 per horizontal rectus and 2.4 ± 0.7 per vertical rectus (n = 14 horizontal and 16 vertical muscles), with some horizontal recti containing more than 20 (Fig. 6C, Fig. S6A-B). We report these numbers for completeness and consistency with the published literature. However, these numbers, as well as previously published palisade-numbers in other species, are absolute counts per muscle that have not been normalized, so differences in count are difficult to interpret across species that differ substantially in eye or EOM size. The structure of an individual ending, by contrast, is independent of these differences and is the feature on which the frontal-versus lateral-eyed distinction was originally drawn^13^.

Rats and tree shrews differed notably in the morphology of their palisade endings. In tree shrews, palisade endings frequently had the form described in frontal-eyed species: the axon reached near the tip of the muscle fiber, sometimes turned back, and gave rise to multiple branches that formed a cuff around the muscle fiber tip, with SYP-positive varicosities both at the tendon level and along the terminal portion of the muscle fiber (Fig. 6F). This elaborate arborization resembles palisade endings in cats and other frontal-eyed mammals^13^. In rats, by contrast, palisade endings were simpler, with fewer branches and varicosities confined to the muscle fiber tip (Fig. 6G), matching the simplified form previously described in this species^13,35^. Additionally, the axons forming palisade endings in rats exhibited synaptic vesicles distributed along the length of the axon immediately preceding the palisade ending (see arrows in Fig. 6G), another aspect of their simplified morphology noted in previous reports. In contrast, synaptophysin labeling in tree shrew palisade endings was concentrated near the terminal portion of the axon, forming larger, more intricate clusters. This distinction was apparent even in relatively simple endings (Fig. 6F–G, right) and was consistent across additional examples from both species (Fig. S6C–D). The more elaborate innervation pattern seen in tree shrews and other frontal-eyed mammals, with clustered synaptic vesicles, may play a role in producing faster or more reliable eye movements.

Together, these results show that, examined with the same methods, tree shrew EOMs contain palisade endings with the morphology characteristic of frontal-eyed species whereas rat EOMs contain the simplified form and mouse EOMs lack them entirely^13^. This anatomical pattern parallels eye-head coupling under head-fixation: eye and head movements are largely uncoupled in tree shrews (Fig. 1), partly coupled in rats (Fig. S1), and, and tightly coupled in mice^5,10^. Determining whether palisade endings are causally related to oculomotor capacity requires additional experiments.

## DISCUSSION

We find that head-fixed tree shrews make saccades frequently and without attempting to move their head (Fig. 1), and that these saccades shift gaze along both the horizontal and vertical axes of the frontal binocular field (Fig. 2). Tree shrews make visually evoked directional saccades (Fig. 3) and, using a gaze-contingent display, we show that they can hold their gaze on a visual target (Fig. 4). This oculomotor repertoire is matched by neural and mechanical specializations: tree shrew visual cortex over-represents the central visual field (Fig. 5), and tree shrew extraocular muscles exhibit elaborate sensorimotor nerve endings similar in morphology to that in frontal-eyed animals with expanded oculomotor repertoire (Fig. 6). Together, these findings show that tree shrews combine lateral eyes and a panoramic field of view with the oculomotor, cortical, and muscular specializations that correlate with eye-driven gaze control in the frontal visual field. Our results do not contradict the established link between lateral eyes and head-driven gaze in the species where it has been studied. Rather, they show that this link is not obligatory. Frontal visual field specialization can be built on a lateral-eyed, panoramic body plan, and eye placement and gaze control are separable dimensions of visual system design.

### Head-independent gaze control in a lateral-eyed mammal

The distinction we draw is not between animals that can and cannot move their eyes. Head-fixed mice make saccades, and can be trained to direct them toward a visual target and to hold gaze near a fixation point^9^. However, mouse saccades are typically accompanied by attempted head movements^5,10^, and even during goal-directed orienting their onset is tightly linked to movement of the head^4^. In freely moving mammals, including mice, rats, tree shrews, and ferrets, saccades can be tightly synchronized with head rotations and followed by compensatory eye rotations that stabilize visual input^16^. Our results show that eye–head coupling during natural locomotion does not preclude independent eye control in tree shrews. Head fixation allows us to test this capacity directly by removing the head’s contribution to gaze. In rats, as previously reported^6^, the eyes move infrequently (Fig. S1A), and most saccades are accompanied by attempted head movements (Fig. S1B–C). In tree shrews, saccade rates are high (median 54 per minute) and the majority (90%) occur with no detectable attempted head movement. Tree shrews therefore can use eye movements, decoupled from the head, for the functions that characterize gaze control in frontal-eyed animals: shifting gaze toward a visual target (Fig. 3) and holding it at a chosen location (Fig. 4). We note that these comparisons span species tested with different tasks and setups, and a direct comparison under identical conditions would strengthen them.

### Eye placement and gaze control as separable dimensions of visual system design

Lateral eye placement, panoramic vision, and a greater reliance on head-driven gaze are often associated across species. Similarly, frontal eyes are associated with greater binocular overlap and more reliance on eye-driven gaze^1^. These correlations are often interpreted in relation to predator-prey ecology: lateral eyes expand visual coverage and help prey monitor predators, whereas frontal eyes increase binocular overlap useful for visually guided predation^7^. Body and head size may further influence whether gaze is redirected primarily by the eyes or the head. These observations can give the impression that eye placement and gaze strategy form a single continuum. But eye placement and gaze strategy need not covary. A recent study across a large range of species shows that even the ecological relationship between predation and eye placement is weaker than traditionally assumed. Once phylogeny is taken into account, eye placement does not differ between predators and other mammals^36^. Our findings reveal a further dissociation: tree shrews retain the panoramic field of lateral-eyed mammals while exhibiting the eye-driven gaze control characteristic of frontal-eyed species. Eye placement determines the extent of visual coverage and binocular overlap, whereas gaze control determines how the eyes are directed within that field.

Tree shrews may not be the only lateral-eyed mammals with this combination of features. Across mammals, the area centralis, a retinal region of high cell density, lies further into the temporal retina as eyes become more lateral^36^, so that it continues to project to the frontal visual field. Thus, many lateral-eyed mammals have a frontally directed retinal specialization despite their panoramic field of view. Whether these animals can redirect that specialization through head-independent eye movements, as tree shrews do, remains largely unknown. Why some lateral-eyed mammals evolved independent control of the eyes while others rely primarily on head movements is also unclear. In tree shrews, one possibility is that the answer lies in their distinctive mode of locomotion.

### What does head-independent gaze control afford an arboreal animal?

We propose that head-independent eye movements resolve a conflict created by acrobatic arboreal locomotion. During leaps, flips, and rotational locomotion the body and head must rotate in the service of locomotion, while vision may need to be directed at a future perch or landing target. Independent control of the eyes could allow an animal to inspect an upcoming target while the head remains engaged in an ongoing movement, potentially favoring ocular gaze control even without frontal eyes. We emphasize that this is a hypothesis, and testing it will require measuring eye and head movements during unrestrained behaviors in which the demands of locomotion and visual sampling may differ. Such measurements would establish whether, and under what behavioral conditions, tree shrews recruit their capacity for head-independent gaze control. If this hypothesis is correct, evolution may favor head-independent eye movements in other lateral-eyed arboreal animals whose locomotion routinely places the head and body in positions that differ from where the eyes need to look. Squirrels, and in particular tree squirrels, are a natural test. They are lateral-eyed, similar in size to tree shrews, and comparably acrobatic, but to our knowledge their eye movements have not been characterized. This view also relates to long-standing debates over whether the visual specializations of primates arose from the demands of arboreal locomotion, visually guided predation, or other aspects of terminal-branch foraging^37,38^.

### Limitations and future directions

Along with head-independent visually guided saccades and fixation, vergence eye movements are another feature of the expanded oculomotor repertoire of primates and other frontal-eyed mammals. Vergence involves coordinated movements of the two eyes in opposite directions, allowing gaze to shift between objects at different distances and supporting binocular depth perception. Given the acrobatic locomotion of tree shrews and the head-independent oculomotor capabilities demonstrated here, we hypothesize that tree shrews may also possess visually guided vergence eye movements. Testing this hypothesis is an important next step which requires a large setup with targets moving in depth. We also did not test smooth pursuit, another hallmark of the primate oculomotor system. Finally, although head fixation allows us to isolate eye movements from the contribution of the head, it precludes the study of gaze during natural behavior. Measuring gaze in freely moving tree shrews during their full range of locomotor behaviors, including leaps and rotations, will be essential for understanding how these oculomotor capabilities are deployed during natural visual behavior.

### A non-primate model for eye-driven gaze

The ability to study head-independent gaze in tree shrews makes it possible to investigate several questions that have historically been studied primarily in primates and cats. These include studying how eye position modulates visual responses, as in the gain fields described in primate cortex^39^; determining how eye position information reaches the brain, whether through proprioceptive signals from the extraocular muscles^40^, or through corollary discharge; and designing virtual reality navigation tasks controlled by eye movements rather than overt locomotion. Tree shrew visual cortex is already well characterized and accessible to modern imaging and viral methods, making these experiments feasible.

## MATERIALS AND METHODS

### Animals and surgical procedures

#### Animals, housing, and husbandry

All procedures were approved by the Cornell University Institutional Animal Care and Use Committee (IACUC) and conducted in accordance with National Institutes of Health guidelines. Eleven adult tree shrews (*Tupaia belangeri*; male and female, 6 months–2 years old) were used. Ten adult rats (Long-evans, all female) were used. We used female rats exclusively as the larger male rats did not fit in our experimental setup. Animals were housed individually or in groups under a 12 h light/dark cycle, with food and water provided daily. Animals undergoing behavioral training were maintained on a regulated water schedule and received juice rewards during experimental sessions. Daily water intake was sufficient to maintain baseline body weight. The data for Figure 5 was collected at the Max Planck Florida Institute for Neuroscience, where the experiments were approved by the Animal Care and Use Committee.

### Headpost implantation

Animals were sedated with midazolam (5 mg/kg, intramuscular [IM]), anesthetized with ketamine (75 mg/kg, IM), and administered atropine (0.5 mg/kg, subcutaneous [SC]) and sustained-release buprenorphine (0.3 mg/kg, SC). The scalp was shaved and disinfected, and local anesthetic (0.3–0.5 mL bupivacaine/lidocaine mixture, SC) was administered at the incision site. Anesthesia was maintained with isoflurane (0.5–2.5%), initially delivered through a nose cone and subsequently through an endotracheal tube with mechanical ventilation (100 breaths/min). Heart rate, ECG, expired CO₂, and body temperature were monitored throughout surgery, with body temperature maintained at 36–37°C. Intravenous fluids (5% dextrose) were delivered through a hindlimb catheter.

Animals were positioned in a stereotaxic frame (Kopf Model 900), and 1-2 metal headplates were secured to the anterior skull over the olfactory bulbs using dental cement (Metabond). The first headplate was used for head-fixation and the second for the IMU. The surrounding skin was sutured closed, and animals recovered for 5–7 days before behavioral acclimation. Surgical procedures for rats were similar, except that rats were not intubated.

### Cranial window implantation for widefield imaging

For widefield calcium imaging, a second surgery was performed at least two weeks after headpost implantation. A metal chamber (10 mm outer diameter, 8 mm inner diameter) was cemented over visual cortex, and a ∼7 mm craniotomy was made, centered approximately 4 mm anterior and 4 mm lateral to lambda. Multiple injections totaling 2–3 μL of AAV9-Syn-GCaMP6s.WPRE.SV40 (10¹³ genome copies/mL; Penn Vector Core or Addgene) were delivered into visual cortex using beveled glass micropipettes (10–20 μm tip diameter) and a Nanoject II injector (Drummond). A total of 3–12 injection sites were made in the cranial window, while the dura remained intact, using pulled glass pipettes. To facilitate spreading, the virus was delivered at two depths, 250 and 500 μm below the cortical surface. After the injections, a durotomy was performed, and a glass coverslip was placed inside the craniotomy and sealed with Vetbond. The skin around the incision was sutured closed. The animals were then placed on a heating blanket in a small cage to recover from anesthesia. The first imaging experiments occurred following a period of 2–3 weeks. The animals continued their acclimation to the recording apparatus during this time. Animals were given twice daily doses of Baytril (5 mg/kg) for 5 days to reduce the -probability of infections.

### Eye and head movement measurements

#### Eye-tracking apparatus

Two infrared cameras (Teledyne FLIR Grasshopper) with macro lenses were positioned so that their imaging planes were parallel to the eyes and their optical axes were roughly aligned with those of the eyes. Each camera was paired with an infrared light source (850 nm) to improve illumination. Frames from the two cameras were captured synchronously at 60 Hz, triggered by a TTL pulse. For the naturalistic video experiment (Fig. 2E, 2F, S2E, S2F), we used a similar camera and IR illumination setup, but with a 100 Hz TTL pulse. To convert eye position from pixels to degrees, we used a two-axis goniometer (Thorlabs GN2) to approximate the eyeball of an adult tree shrew. A custom-designed pole was attached to the goniometer stage so that its tip could rotate about two axes with a radius of 3.9 mm^41^. The goniometer was placed on the stereotax so that the position of the tip relative to the camera approximated that of the eye relative to the camera. The goniometer was rotated from −15° to +15° horizontally and from −10° to +10° vertically. We recorded the position of the tip in pixel space (*p*_1_, *p*_2_) at two arbitrary angles (*θ*_1_, *θ*_2_). The pixel to degree conversion ratio (r) was calculated as 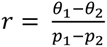. This measurement was then cross-validated by randomly selecting two positions in pixel space, calculating the corresponding angular displacement using the ratio, and comparing it with the angular displacement of the goniometer (ground truth). This procedure was repeated for both eyes.

### Real-time data acquisition and eye-tracking

We used Bonsai-Rx^42^ for real-time processing of the camera streams and to control peripherals such as speakers, monitors, and Arduino Uno boards. The two cameras were synchronized by a custom TTL signal generated by an Arduino. Juice was delivered with a programmable syringe pump (New Era Pump Systems, NE-1000), and the timing of delivery was controlled by a pulse generated by a second Arduino. We used DeepLabCut^43,44^ to train an eye-tracking model that tracks 8 points around the pupil and 4 points around the eye. To build the model, we first cropped the videos to show only the eye, then manually labeled all points on 100–150 frames. We periodically added newly labeled frames to the training set to maintain model performance, and the model output was also verified visually. For online eye tracking, we used DeepLabCut-Live^45^, through its plugin designed specifically for Bonsai-Rx. The online model inferred the point locations for every frame in the stream. To ensure real-time performance, the camera streams were downsampled by a factor of 3. The 8 inferred points around the pupil were then fitted with an ellipse using OpenCV, and the center of the ellipse was taken as the center of the pupil. Of the 4 points around the eye, only the anterior and posterior points were used to calculate the center of the eye. Because these points were expected to remain largely stable throughout the experiment, they were first Gaussian-filtered (σ=3) to remove any imaging jitter. Finally, eye movement per frame was calculated by subtracting the pupil center from the eye center.

### Measurement of attempted head movements

We used a 9-DoF IMU (SparkFun Electronics, ICM-20948) to measure head acceleration as a proxy for the force generated by the animal during attempted head movements. Only the three accelerometer axes (*a_x_*, *a_y_*, *a_z_*) were used for analysis, and the accelerometer sensitivity was set to ±2 g. The IMU was connected to an Arduino via the serial peripheral interface (SPI) and programmed to stream data continuously. However, the Arduino sent a data packet (containing all 9 axes of data and a checksum) to Bonsai-Rx only upon receiving a TTL pulse, which was detected by an interrupt service routine. This ensured that the IMU packets were synchronized with the camera frames. We verified the temporal fidelity of the IMU signal manually by applying a periodic vibration and confirming that the output was sinusoidal at the same frequency.

### Force-sensor validation of IMU measurements

To validate IMU measurements of attempted head movements, we used a force-sensor apparatus adapted from Hughes et al.¹⁷ (Fig. S1D). Signals from three orthogonal axes were acquired through the analog inputs of an Arduino Uno and synchronized with the eye-tracking cameras and IMU using TTL pulses. Data were analyzed from one session in one tree shrew, using the first 70% of the recording after excluding its initial 1%. Saccades were detected automatically from horizontal eye position. IMU signals were band-pass filtered at 3–15 Hz and force-sensor signals at 2–12 Hz. For each signal, we calculated the Hilbert envelope of each axis and combined the three axes using their Euclidean norm. For each saccade, we extracted the peak of each combined envelope within ±0.25 s of saccade onset. The relationship between peak force-sensor signal (mV) and peak IMU acceleration (g) was quantified using Pearson’s correlation coefficient and a least-squares linear fit.

### Offline saccade detection and analysis

For the offline analysis, we processed the eye videos from each session using a custom Python script that integrates DeepLabCut. We only kept frames where the average confidence of the pupil points was at least 0.95 and the average confidence of the eye points was at least 0.9, and replaced all other frames with NaN. We used OpenCV to fit an ellipse to the 8 points around the pupil. We used the anterior and posterior points around the eye to calculate the eye center. We also tracked the distance between these two points, and if it deviated from the mean by more than two standard deviations, we discarded those values, since this would violate the rigid-body constraint of the eye. We used the dorsal and ventral points to detect blinks. To do that, we tracked the distance between these two points, and if its rate of change crossed a velocity threshold (99.99th percentile), we considered that frame a blink. Since this only captures the frames when the eye is closing or opening, we applied binary dilation to the blink frames so that the frames when the eye stays closed are also counted as blinks. We replaced all blink frames with NaN and later used linear interpolation to fill in all the NaN frames. Then, we obtained the eye-centered pupil position by subtracting the eye center from the pupil center, and median-filtered it with a window size of 3 frames to remove any jitter. Finally, we convert the eye-centered pupil position from pixel to degrees by the conversion ratio explained in the earlier section. We get eye movements in yaw and roll directions using this method. To obtain the pitch eye movements, we directly use the angle of the fitted ellipse.

For saccade detection, we analyzed each axis of eye movements separately. We used a custom designed velocity threshold-based saccade detection algorithm ^46^. The velocity threshold is iteratively estimated based on the median absolute deviation of the eye movement velocity distribution. We further filtered the candidate saccades based on a set of criteria: a) duration between 2 samples to 80 ms b) minimum amplitude = 1 ° c) difference between successive samples to be at least 80 ms d) only one velocity peak during the saccade. We manually verify all saccades in rats and 1098 randomly sampled saccades in tree shrew. More details regarding the manual verification are provided in the next section. Although we detected saccades in each axis separately, to find out how much they overlap across the three axes, we pooled the eye velocity time series across sessions for each axis and plotted their covariance matrix.

Every automatically detected saccade in the rat sessions (590 in 18 sessions) and a random sample of tree shrew detections (1,098 in 76 sessions) was inspected visually and classified as a saccade or not. Inspection confirmed 1,091 of the 1,098 tree shrew detections (99.4%) and 337 of the 590 rat detections (57%). Validation rates were lower in rats due to increased head, face, and whisker movements. So that both species were analyzed with the same rule, analyses of individual saccades (main sequence, saccade-triggered averages, and head acceleration around each saccade) used only saccades confirmed by inspection, at most 1,000 per species drawn at random: 1,000 of the 1,091 in tree shrews and all 337 in rats; sample sizes therefore differ by design.

For the main sequence analysis, we fitted the peak speed of the saccade (*v_p_*) and the amplitude (*a_s_*) as per the following power-law equation: 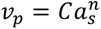, where *C* is the coefficient and *n* is the exponent.

### Offline detection of attempted head-movements

To detect attempted head movements, we first band-pass filtered the accelerometer signal in each axis between 5 and 20 Hz using a 4th-order zero-phase Butterworth filter. We chose these band-pass parameters by comparing the frequency spectrum of the accelerometer signal in each axis between the experimental sessions and a recording with no animal on the rig. We then used the Hilbert transform to get the envelope of the filtered signal in each axis and combined the three axes by calculating the Euclidean norm of the envelopes. To express the envelope in noise units, we used the standard deviation of the unfiltered accelerometer signal recorded with no animal on the rig, with the three axes combined (0.00127 g), as one unit. An attempted head movement is considered whenever the normed envelope was above twice standard deviation. A saccade was categorized as "with head movement" if the normed envelope crossed this threshold within ±0.5 s of saccade onset, and as "without head movement" otherwise. Here and in saccade-triggered analyses, saccade onset refers to the time of the saccade’s peak velocity. 95% confidence intervals on the percentage of saccades in each category were calculated with the Wilson score method from the number of saccades in the category and the total number of saccades, pooled across animals and treating saccades as independent. Unsigned saccade amplitude was compared between the two categories with a two-sided Mann-Whitney U test.

### Behavioral acclimation & training

#### Habituation to head fixation

Animals underwent 1–3 weeks of acclimation to handling, head fixation, and the recording apparatus. During recording, the implanted headplate was secured to a custom-built stereotaxic holder, and body movement was constrained by a transparent acrylic tube. Head-fixation duration was gradually increased, with juice rewards used to reinforce calm behavior. Animals were maintained on a regulated water schedule during acclimation and training (typically 15–30 mL/day) and received juice rewards during the sessions.

### Saccade-to-target training

Animals were trained to make visually evoked directional saccades through progressively more demanding reward contingencies. Training began by rewarding any detectable eye movement, followed by saccades made within 1-2 s of visual cue onset. Rewards were then restricted to saccades directed toward a visual target appearing on the left or right side of the monitor, and the range of rewarded directions was progressively narrowed. Animals typically spent 2–5 sessions at each stage, advancing when performance reached approximately 75%.

### Fixation training

Fixation training proceeded in two stages. Animals were first trained to maintain a stable eye position following a visual cue, with rewards contingent on the absence of saccades for progressively longer durations (300–750 ms). Once animals reliably maintained fixation, they were introduced to a gaze-contingent fixation task in which estimated eye position was displayed in real time as a movable marker on the monitor. Animals learned to use this visual feedback to bring the marker onto a target and maintain its position there for a specified duration. Training began with large targets that were progressively reduced in size as performance improved. In the final task, targets appeared at one of two positions, and animals were rewarded for maintaining the gaze-contingent marker within the target for 500 ms. Trials terminated without reward if the fixation criterion was not met within 11 s.

### Behavioral tasks and related analysis

#### Visual stimulus presentation

We used BonVision^47^ to generate and present visual stimuli on a monitor. The luminance and contrast of the monitor were: white 79.9 cd/m² (CIE 1931 x = 0.304, y = 0.331; CCT ≈ 6968 K), black 1.20 (95% CI 0.77–1.64) cd/m², Michelson contrast 0.970 (95% CI 0.960–0.981). The monitor had a resolution of 1920 × 1080 pixels and was placed 47 cm from the eyes, spanning a visual field of 54° × 32°. The center of the monitor was aligned with the midpoint between the eyes.

### Saccade-to-target task

Each trial began with a central blue cue (∼3° diameter) presented for 0.7 s, followed by a visual target appearing randomly to the left or right, approximately 16° from the monitor center. Targets had a radius of 6–9° and contained a drifting grating (1–2°/s) to increase salience. The animal was rewarded with 0.02 mL of diluted apple juice (1:1 with water) if its first saccade occurred within 1 s of target onset and was directed toward the target. Saccade direction was calculated from the angle of the eye-displacement vector relative to the target direction, with deviations of ≤90° scored as correct. Trials were classified as valid if a saccade occurred within 1 s of target onset and invalid otherwise. Valid trials were scored as correct or incorrect according to the direction of the first saccade. Trials ended after the first saccade or at the 1 s timeout, followed by a 4 s inter-trial interval.

Validity was calculated as the fraction of all trials containing a saccade within 1 s of target onset, and accuracy as the fraction of valid trials in which the first saccade was directed toward the target. To examine the temporal evolution of directional accuracy, we calculated the fraction of correct first saccades in sliding 0.3 s latency windows, centered at 0.05 s intervals from 0 to 1.0 s. Pointwise 95% Wilson score confidence intervals were calculated from the number of correct saccades and the total number of saccades in each window. For a pre-cue control, we identified the last saccade, if any, within the 1 s preceding target onset on each trial and classified its direction relative to that trial’s upcoming target using the same ±90° criterion

### Cued fixation task

In this task, each trial starts with a fixation cue with a radius of ∼12°. The animal has to fixate for 400–700 ms. If the animal does not fixate, the task does not move forward until a 11 s timeout, after which the next trial starts after a 4 s delay. If the animal fixates successfully, it receives a 0.02 ml drop of the juice mixture as a reward.

### Gaze-contingent fixation-on-target task

In this task, a target of similar size to the one in the cued fixation task appears randomly on the left or right after a blue cue. The animal is expected to use feedback from the gaze-contingent marker to move it onto the target and hold its gaze there for 400–700 ms. As in the cued fixation task, there is a 11 s timeout, and the animal receives the same reward upon successfully completing the trial.

We obtained the real-time eye position using DeepLabCut-Live in Bonsai-Rx, as described above. We used these positions to roughly estimate where on the 2D plane of the monitor the animal was looking (*x*, *y*), using a simple linear transformation and manual calibration:

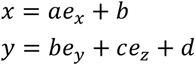

We estimated the values of the coefficients by presenting a very salient stimulus on the monitor. We used BonVision within Bonsai-Rx to show the animal its estimated gaze position on the monitor by creating gaze-contingent marker of ∼4° diameter at the *x*, *y* position.

### 3D eye-in-head models

#### Geometric modeling of eye movements and gaze

To relate eye rotations to pupil motion seen by the eye-tracking cameras and to gaze-shifts in the visual field, we built three-dimensional models of the eyes in the skull of a tree shrew and a macaque in Blender, scripted in Python through the Blender API. The tree shrew skull was a smoothed surface mesh of a *Tupaia belangeri* skull scanned in the lab. The macaque skull was the "Macaque Cranium" model by the UNCG Imaging Lab (https://skfb.ly/O8YY), used under a CCBY 4.0 license. Skulls were scaled to a common width, and models are therefore in arbitrary units. The models were used to illustrate the geometry and to derive qualitative predictions about which rotations appear on camera as pupil translation or as pupil torsion and which shift gaze. They were not used to convert pupil position into gaze angle.

### Retinotopic mapping and cortical surface area analysis

#### Widefield calcium imaging

Retinotopic maps of primary visual cortex (V1) were obtained using widefield calcium imaging of GCaMP6s-expressing neurons in awake, head-fixed tree shrews, using previously established methods^27^. A large cranial window (10mm chamber, 7 mm visible cortex) was implanted over visual cortex, and multiple injections of a viral vector containing GCaMP6s were made within the exposed cortex. Widefield imaging commenced following a 3 week period to allow expression of the virus. Once acclimated to the head-fixation apparatus, animals passively viewed visual stimuli presented on a monitor positioned in the frontal visual field. Imaging was performed using a Bergamo II microscope (Thorlabs) with blue LED excitation and a Zyla 5.5 sCMOS camera (Andor) controlled by μManager. GCaMP6s fluorescence was recorded at approximately 15 Hz, with images acquired at 640 × 540 pixel resolution. Excitation power measured at the objective ranged from 0.2 to 0.8 mW using 2× or 4× air objectives (Olympus).

To map azimuth and elevation, a bar swept horizontally and vertically across the monitor, respectively (Fig. 5A). Visual stimuli were generated using PsychoPy and presented on a monitor positioned in the frontal visual field. Bars were 3° wide and moved at 4°/s, with 30–50 repetitions per condition. The monitor had a resolution of 1920 × 1080 pixels and was positioned approximately 20 cm from the animal, subtending approximately 120° × 70° of visual space. Retinotopic maps were generated using the phase of the stimulus-evoked fluorescence response at each cortical location, following established temporally encoded mapping approaches^24^. The resulting phase maps were used to assign azimuth and elevation coordinates to cortical locations within the imaging window (Fig. 5B). Imaging data were processed using custom MATLAB and Python scripts. Images from individual recording sessions were first registered to a reference image of cortical blood vessels acquired during the initial imaging session, followed by more precise offline registration using pixel correlations. Vascular artifacts were removed from the retinotopic maps before identifying the representations of the vertical and horizontal meridians and measuring the cortical area corresponding to the central visual field (Fig. 5B). The vertical meridian was identified from the mirror symmetry of azimuth representations across the two hemispheres, using the median phase of corresponding cortical locations. The horizontal meridian was assigned to the center of the stimulus monitor, accounting for a 300 ms fluorescence response delay. Phase maps were smoothed independently for each hemisphere with a Gaussian filter (σ = 12 pixels). Pixels with trial-to-trial response coherence below 0.5 and those within 14 pixels of the cortical midline were excluded.

### Reconstruction of the tree shrew cortical flatmap

We generated a cortical flatmap from the 3D reconstruction of the tree shrew brain provided in^21^, derived from 9.4 T anatomical MRI scans. Using FreeSurfer^48^, Pycortex^49^, and Blender, we inflated the cortical surface, removed medial portions of the corpus callosum, and introduced five cuts to flatten the cortex while minimizing distortion. V1 boundaries were identified using cortical curvature and published anatomical and retinotopic maps. Because the dorsal V1 boundary was not clearly resolved in the atlas surface, we reconstructed a higher-resolution curvature map from MRI scans of two additional tree shrews. We traced the dorsal boundary using SUMA/AFNI^50^, registered it to the original flatmap, and verified it against ^25^. The medial and caudal boundaries were traced using the map of Kaas (1980). Total V1 surface area was calculated using Green’s theorem, yielding 62.5 mm², consistent with the published estimate of 66 mm²^28^.

### Quantification of central visual field over-representation

The cortical area representing the central visual field was calculated by counting pixels within the specified retinotopic boundaries and converting pixel counts to mm² using known scaling. To account for incomplete coverage of the central visual field, we divided visual space into 0.5° × 0.5° bins and identified bins represented by the selected cortical pixels, following previously published approach^23^. Gaps between occupied bins were closed using morphological binary closing (15 iterations). The resulting visual field area was calculated from the number of covered bins. Total visual field coverage of V1 was calculated as the area of a polygon connecting the outermost receptive-field locations reported by Kaas et al^26^. Cortical and visual-field fractions were calculated separately for each hemisphere and averaged before computing the over-representation ratio.

### Validation of tree shrew ratio

We digitized the V1 boundary and retinotopic grid from Bosking et al.^25^ (Fig. 10A). The region bounded by the relevant azimuth and elevation contours was identified using polygon geometry to calculate its surface area. To compare the published map with the portion sampled by our imaging window, we generated a triangular mesh of the digitized region (0.055 mm spacing) and interpolated azimuth and elevation across the mesh using a cotangent-Laplacian interpolation constrained by the traced retinotopic contours. We then summed the areas of mesh triangles whose visual field coordinates fell within the same 0.5° bins represented in our widefield maps. An independent estimate of over-representation was also calculated using the full cortical region in the published map and the V1 surface area reported in Sesma et al.^28^.

### Cross-species comparisons

For squirrel monkeys, we used the cortical surface areas between iso-eccentricity contours reported in^22^. Cumulative cortical area was interpolated linearly against log eccentricity. The area representing the central region was calculated by integrating across eccentricity. Total V1 area included the monocular crescent, whose visual-field extent was estimated using cortical magnification in the outermost measured eccentricity band. For mice, we analyzed the published azimuth and altitude imaging-based maps^23^. V1 was segmented using the visual field sign and morphological filtering procedures described in that study. The visual-field origin was defined at the junction of V1, LM, and RL, and cortical area was measured by counting pixels within the retinotopic boundaries. Visual-field coverage was calculated using the same 0.5° binning procedure described above. We independently analyzed the electrophysiological map of Wang and Burkhalter ^29^ by digitizing the locations and receptive-field coordinates of 19 recording sites and interpolating retinotopy across V1 using thin-plate splines. Uncertainty in this estimate was assessed using 200 random perturbations of the digitized coordinates (SD = 0.06 mm for recording locations and 3° for receptive-field coordinates), with the 5th–95th percentile range reported. For the primary cross-species comparisons, uncertainty in the reference positions was assessed using the ±5° shifts described in the Results.

### Extraocular muscle anatomy and histology

#### Tissue extraction and fixation

Extraocular muscles (EOMs) were obtained from 4 tree shrews and 7 rats. Animals were deeply anesthetized, euthanized via barbiturate overdose, and intracardially perfused with physiological saline (0.9% NaCl in PBS) followed by 4% paraformaldehyde in PBS. The eyes, with the EOMs attached, were then removed from the orbits and postfixed for 2 hours before being transferred to PBS containing 0.05% sodium azide and stored at 4°C until further processing. The four rectus muscles (medial, lateral, superior and inferior) were carefully dissected away from the eye leaving as much of the myotendinous junction as possible. Muscles were then stained via immunofluorescence and mounted on slides for imaging.

### Immunofluorescence labeling

To label palisade endings, extraocular muscles were stained with antibodies against neurofilament and synaptophysin, as well as with phalloidin. Neurofilament was used to label axons, synaptophysin to label synaptic vesicles, and phalloidin, a toxin which binds to actin, to label muscle fibers.

To prepare the tissue for staining, EOMs were first washed 3 times for 5 minutes each in PBS containing 1% Triton (PBS-T). Tissue was then blocked for 2 hours with 10% normal donkey serum (Fisher Scientific, OB003001) in PBS-T. Tissue was then incubated in all primary antibodies for 48 hours, washed with PBS-T again, and incubated in all secondary antibodies and phalloidin for 24 hours. Finally, tissue was washed again and transferred to PBS with 0.05% sodium azide for storage at 4°C until being mounted on slides. All washing and incubation was done on an orbital shaker at room temperature. Staining protocols were based on published methods^13^. The antibodies and stains, along with the concentration used and their supplier and catalog number are listed in the following table.

|  | **Antibody/stain** | **Concentration** | **Supplier (catalog number)** |
|---|---|---|---|
| **Primary antibodies** | Chicken anti-neurofilament | 1:2500 | Fisher Scientific (AB5735MI) |
|  | Rabbit anti-synaptophysin | 1:300 | Fisher Scientific (AB9272MI) |
| **Secondary antibodies** | Donkey anti-chicken Alexa Fluor 647 | 1:500 | Fisher Scientific (A78949) |
|  | Donkey anti-rabbit Alexa Fluor 555 | 1:500 | Fisher Scientific (A31572) |
| **Stains** | Phalloidin (Alexa Fluor 488) | 1:200 | Fisher Scientific (A12379) |

For mounting on slides, in both species, the EOMs were cut transversally into two pieces and only the distal part of the muscle containing the myotendinous junction was mounted. Due to the large, thick and irregular shape of the tree shrew EOMs (particularly the medial and sometimes inferior rectus muscles), this was necessary to flatten the muscles enough to obtain good confocal images. However, in the rat, muscles were thinner and the entire muscle could also be mounted without cutting. Muscles were whole-mounted in Fluoromount-G and stored in the dark at 4°C until imaging.

### Confocal microscopy

Fluorescently labeled EOMs were imaged using a confocal laser scanning microscope (LSM 710 and LSM 980, Carl Zeiss). For the LSM 710, images were captured using a 10x air objective with 3x digital zoom or a 40x water immersion objective with a z-step size of 3 or 4.29 µm respectively. For the LSM 980, images were captured using a 40x objective with a z-step size of 2.5 µm. The entire myotendinous region was imaged and optical sections were collected through the entire thickness of the muscle. Images were acquired using lasers with excitation wavelengths of 488, 561, and 633 (LSM 710) or 639 (LSM 980) nm. Close up images of individual palisade endings (Fig. 6 F,G) were acquired using a LSM 980 with a 63x oil immersion objective and a z-step size of 0.25-1.5 µm. Image post-processing was performed using FIJI. Brightness and contrast were enhanced as required, and maximum intensity projections were generated for visualization.

### Palisade ending identification

The criteria used for identifying a PE was as follows: an axon coming from the muscle and ending near the tendon, making at least one contact with the end of a muscle fiber, with synaptophysin present within 50µm of the fiber end. Muscle fibers that ended more than 150µm before the tendon were not included. Muscle fiber ends were defined as the point in which the fiber tapers to ½ of its original width. Due to the difficulty of identifying a 3D u-turn structure from 2D images, palisade endings that did not clearly exhibit a u-turn of the axon were still included. Counts were then reviewed by 2-3 independent reviewers before being finalized.

## ACKNOWLEDGEMENTS

We thank Mason Raffo, Zed Deas, and Sharvari Saraf for assistance with experiments and Haley Goroway for assistance with animal care. We thank David Fitzpatrick, Susan Freling, Nicole Shultz and Idris El-Amin at the Max Planck Florida Institute for Neuroscience for support in establishing our tree shrew colony. We thank Erica Feldman, Trevor Toman and the Cornell CARE team for their support in ensuring animal welfare. Imaging data was acquired through the Cornell Institute of Biotechnology’s Imaging Facility (RRID:SCR_021741) with NIH funding for the shared Zeiss LSM 710 (1S10RR025502) and 980 (S10OD036283) confocal microscopes.

## AUTHOR CONTRIBUTIONS

Conceptualization (RP, EK, MSS); Methodology (RP, EK, RA, MSS); Investigation (RP, EK, RA, ET, AB, MSS); Formal analysis (RP, EK, MSS); Visualization (RP, EK, ET, AB, MSS); Writing – original draft (MSS); Writing – review & editing (RP, EK, ET, AB); Supervision (MSS); Funding acquisition (MSS).

## AI DISCLOSURE

Generative AI tools (Claude, Claude Code, ChatGPT, GitHub Copilot) were used to assist with writing and debugging analysis and visualization code, converting code from MATLAB to Python, and to improve the text of the manuscript. All AI-assisted recommendations were reviewed, tested, and validated by the authors. All analyses, results, and interpretations were performed and verified by the authors. The authors take full responsibility for the content of the manuscript.

## DATA AND CODE AVAILABILITY

Analysis code to reproduce the figures in this paper will be made available at https://github.com/SarvestaniLab/2026tsgazecontrol. Data will be made available at request.

## SUPPLEMENTARY DATA

**Figure S1:**
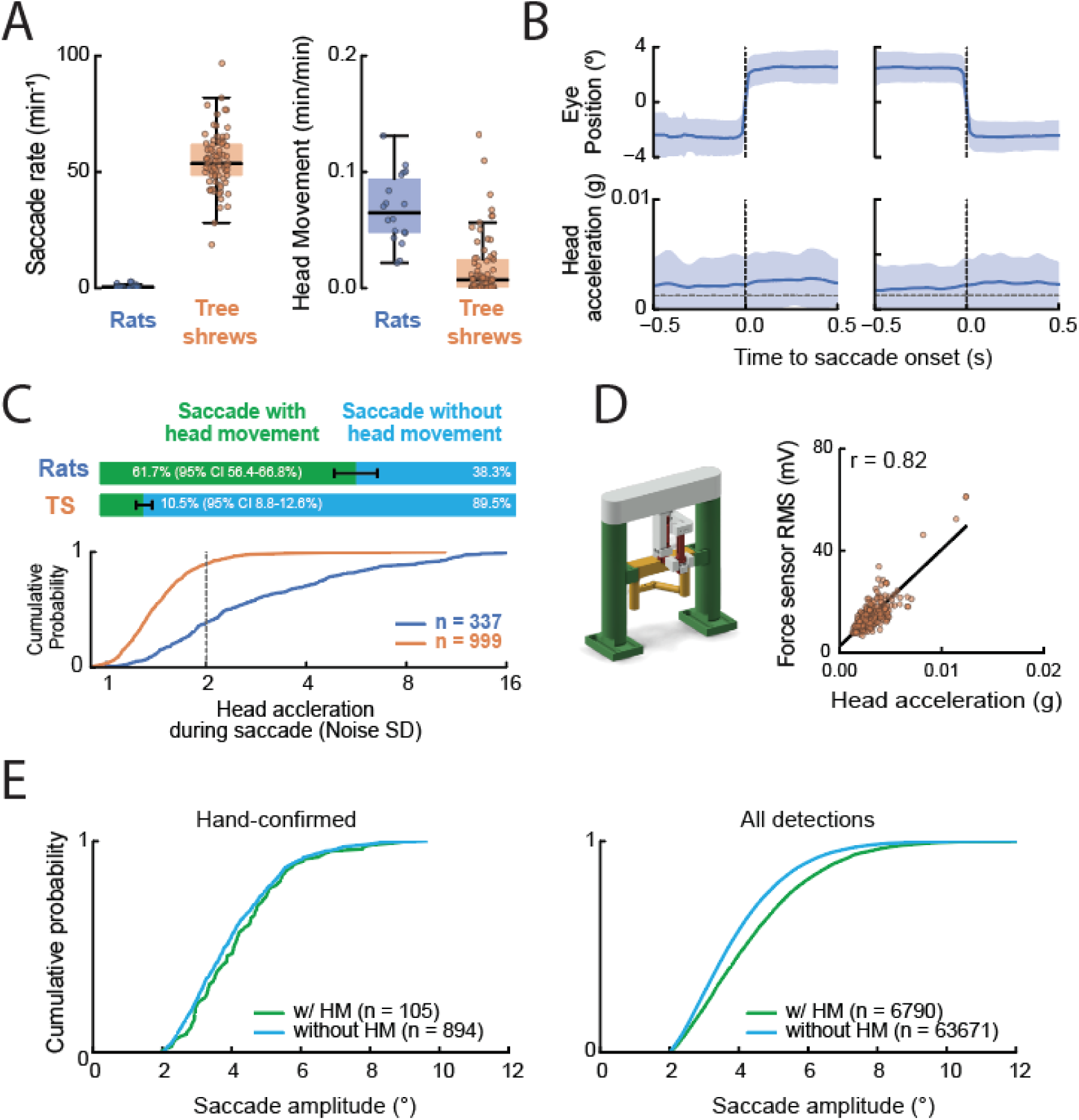
Supporting data for Figure 1. **A)** Saccade rates (left) and time spent making attempted head movement (right, minutes with head acceleration above threshold per minute of recording) in head-fixed rats and tree shrews. 337 saccades were visually verified in 2 rats; 1000 saccades were verified in 2 tree shrews. Animals were passively viewing visual stimuli. Dots, sessions (18 in rats, 76 in tree shrews); boxes, median and interquartile range. Rats made a median of 0.51 saccades per minute, compared with 53.7 in tree shrews, and attempted head movements for 0.065 min per min, compared with 0.007 in tree shrews. Saccade rates are corrected for the fraction of detections confirmed by visual inspection (Methods). **B)** Saccade-triggered average of yaw eye position and head acceleration during saccades in rats. The dashed line is the SD of the unfiltered no-animal signal (0.00127 g). **C)** Top: percentage of saccades with and without attempted head movement in rats (61.7% [95% CI 56.4–66.8%] and 38.3%; n = 337) and tree shrews (10.5% [95% CI 8.8–12.6%] and 89.5%; n = 999); error bars, Wilson 95% CI. Bottom: cumulative distribution of peak head acceleration within ±0.5 s of each saccade, in units of the no-animal SD; the dashed line is the head-movement threshold (2 SD). **D)** Left: force-sensor apparatus used to validate the head-mounted IMU. Right: relationship between peak force-sensor signal and peak IMU-measured head acceleration during individual saccades (n = 245 hand-confirmed saccades). Line, least-squares fit; Pearson’s r = 0.82. **E)** Cumulative distribution of tree shrew saccade amplitude for saccades with and without attempted head movement. Left: confirmed saccades (median 4.1° with, 3.8° without; p = 0.12). Right: all detections (median 4.2° with, 3.7° without; p < 10⁻¹⁰⁰). Two-sided Mann-Whitney U test.

**Figure S2:**
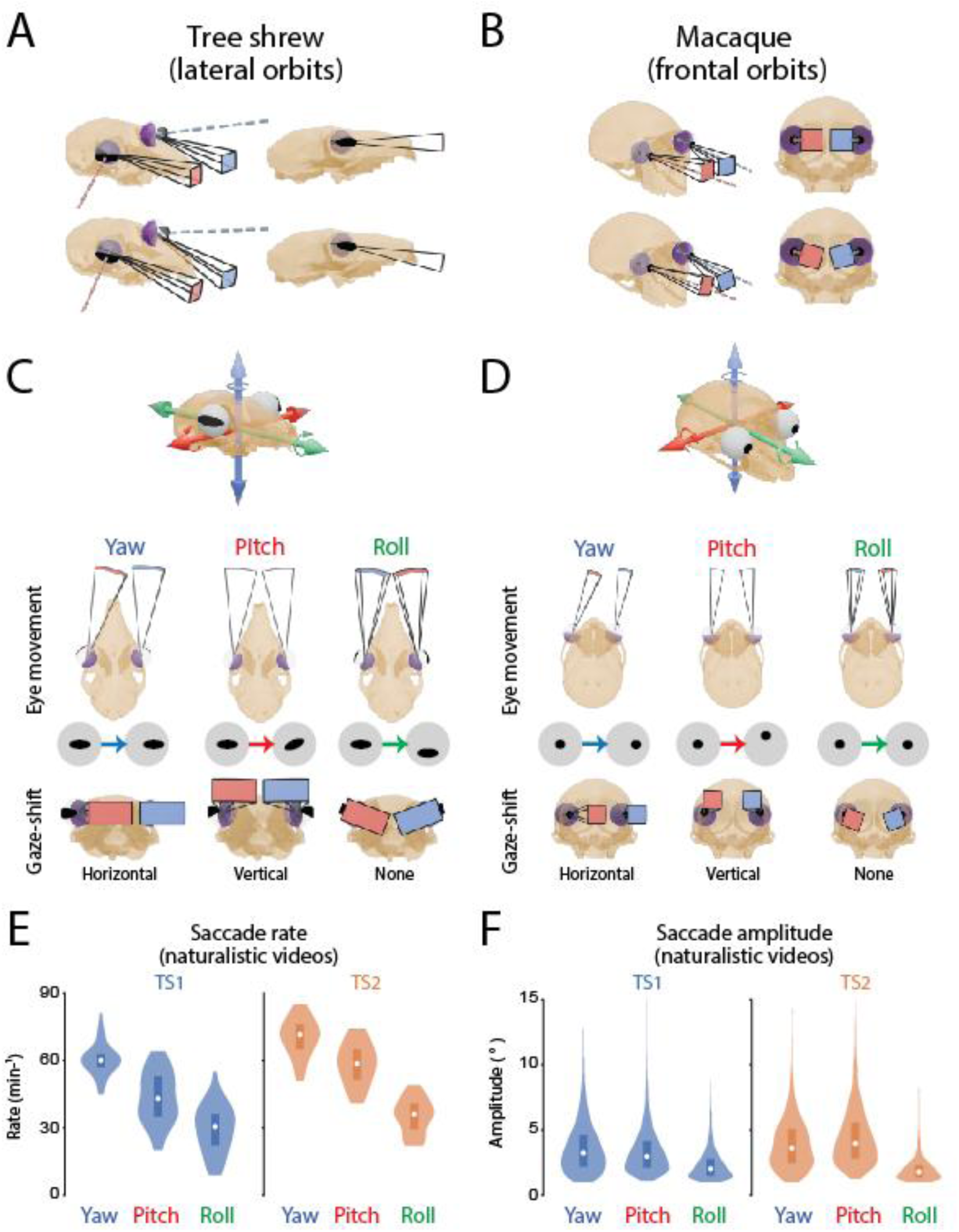
Supporting data for Figure 2. A) In lateral eyed animals such as tree shrews, torsional movement of the eyes, defined as rotations along the optical axis (see Figure 2), produce vertical gaze-shifts. B) In frontal-eyed species such as primates, the same torsional movement produces no shift in gaze, and rather a rotation of the image. C-D) Orbit position influences relationship between pupil displacement visible on a camera placed parallel to the eye (perpendicular to optical axis) and the consequence of that eye movement on gaze-shifts in the visual field. Yaw rotations look like a horizontal displacement of the pupil for both lateral and frontal eyed species and produce horizontal gaze-shifts. Pitch movements look like torsional/angular rotation of the pupil in lateral and vertical displacement of the pupil in frontal eyed animals, although the consequence is a vertical gaze-shift in both species. Roll movements produce a vertical displacement of the pupil in lateral and a torsional/angular rotation of the pupil in frontal eyed animals, and the consequence is a rotation of the retinal image instead of a gaze-shift in both species. E-F) Individual animal data for Saccade rate (left) and amplitude (right) for the three types of eye-movements, recorded while animals (n=2) passively viewed naturalistic movies.

**Figure S3:**
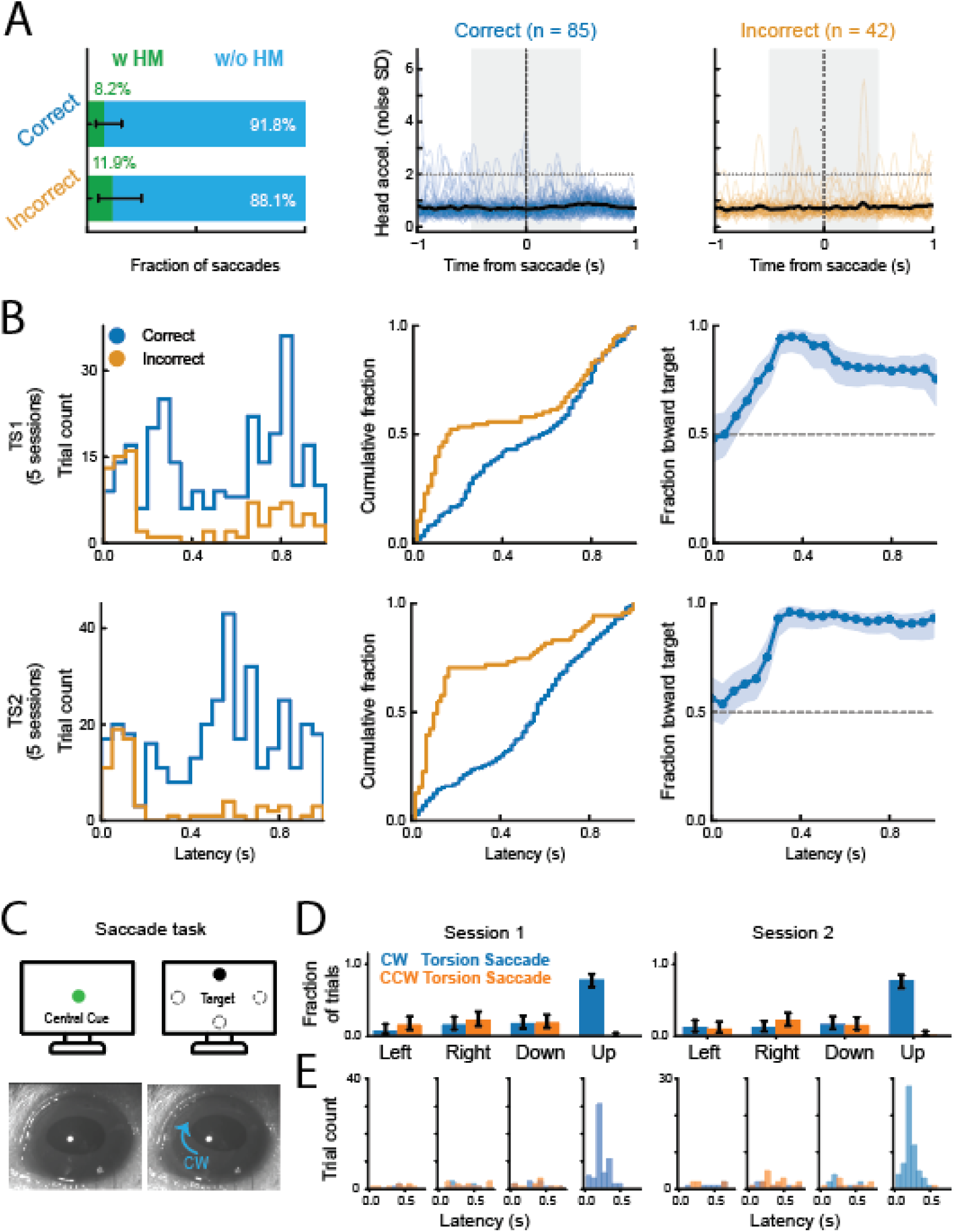
Supporting data for Figure 3. **A)** Attempted head movement around visually evoked saccades in TS1, pooled across three saccade-to-target sessions recorded with the head-mounted IMU. Left: fraction of correct (n = 85) and incorrect (n = 42) first saccades with (green) and without (blue) attempted head movement. As in Figure 1, a saccade is classified as with head movement if the head-acceleration envelope exceeds 2 SD of the no-animal signal within ±0.5 s of saccade onset. Error bars show 95% Wilson CIs of the fraction with head movement (correct 8.2%, incorrect 11.9%). Middle and right: head acceleration around each correct (middle) and incorrect (right) saccade. Thin lines are individual saccades, and the thick black line is the mean. Grey shading marks the ±0.5 s classification window, and the dotted line the head-movement threshold. Individual traces crossing the threshold show that the IMU detects attempted head movements during the task. These movements were not aligned to saccade onset, and mean head acceleration stayed near the noise floor around both correct and incorrect saccades. The small rise in the mean about 0.5 s after correct saccades reflects licking during juice delivery. **B)** Saccade-to-target performance for TS1 (top; n = 379 valid trials) and TS2 (bottom; n = 422 valid trials), pooled across five sessions per animal. Left: distributions of first-saccade latencies for correct (blue) and incorrect (orange) trials. Middle: cumulative latency distributions of correct and incorrect saccades. Right: fraction of correct first saccades in sliding latency windows (0.3 s wide, 0.05 s steps). Shading shows pointwise 95% Wilson CIs; the dashed line shows 0.5. **C)** Task with vertically displaced targets. A central cue (green) is followed by a target (black) appearing randomly at one of four positions (left, right, up or down; dashed circles mark the possible target locations). Bottom: eye-tracking camera view showing clockwise (CW) pupil torsion associated with upward pitch saccades (blue arrow; see Fig. 2D). **D)** Fraction of trials containing CW (blue) or counterclockwise (CCW, orange) torsion saccades for each target position in two example sessions. Error bars show 95% Wilson confidence intervals from the trial counts in each session. E) Distributions of torsion-saccade latencies relative to target onset for the sessions in D.

Although our saccade task focused on horizontal saccades (Fig. 3), we also wondered whether tree shrews could make saccades to targets displaced along the vertical axis of the visual field. This would require animals to make pitch saccades, as predicted from our eye-in-head model (Fig. 2), which produce little translational pupil displacement on camera but are instead visible as torsion of the elliptical pupil (Fig. 2D). The task used here was similar to that for horizontal saccades, except that in addition to left and right targets, targets also appeared randomly above (Up) and below (Down) the center of the monitor. Only one target appeared at a time (Fig. S3C). Upward targets reliably evoked clockwise (CW) torsion saccades—the camera signature of an upward pitch rotation—on the majority of trials, at short latencies (<0.5 seconds) after target onset, whereas counterclockwise (CCW) torsion saccades on these trials were nearly absent (Fig. S3D–E). Although CW saccades also appeared during left and right target trials, they were much less frequent than during upward-target trials. Tree shrews therefore direct visually evoked saccades along both axes of the frontal binocular field. Notably, downward targets did not evoke the complementary CCW (downward pitch) torsion saccades: torsion-saccade rates on downward-target trials remained at the low baseline seen on horizontal-target trials (Fig. S3D). A likely explanation is that the tree shrew’s long snout occludes a region of the lower frontal visual field, so the stimulus may simply not have been visible.

**Figure S4:**
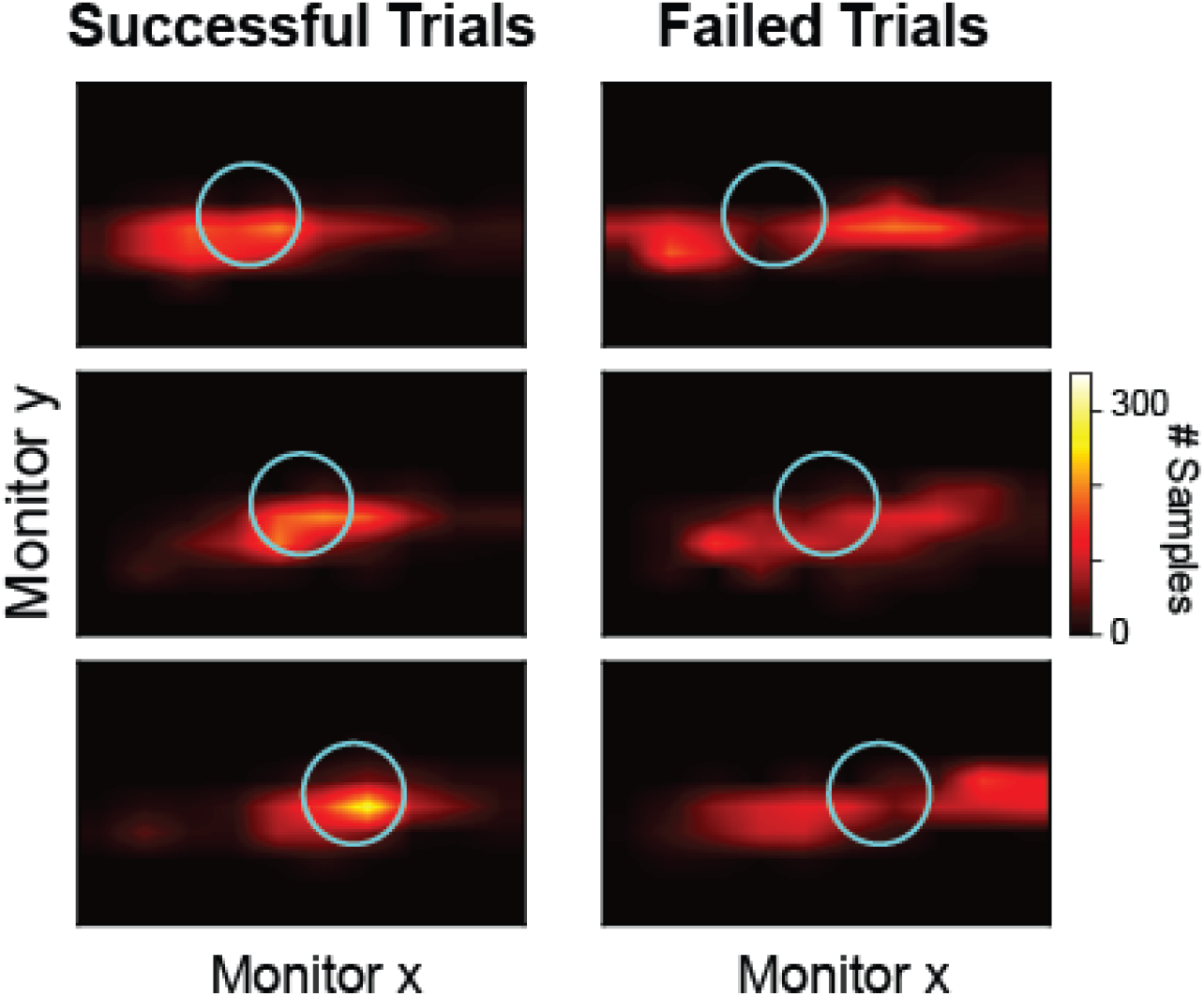
Supporting data for Figure 4. **Gaze density tracks target location and size.** Heatmaps of gaze position on the monitor for three sessions with a single fixation target shown at different monitor positions (rows). Left column: gaze density across successful trials where the animal was able to bring the gaze-contingent marker onto the target. Right column: failed trials only. In all cases, gaze density is concentrated around the target and follows it as its position and size vary, including on trials that ended in failure, indicating that animals directed gaze toward the target rather than encountering it by chance.

**Figure S5:**
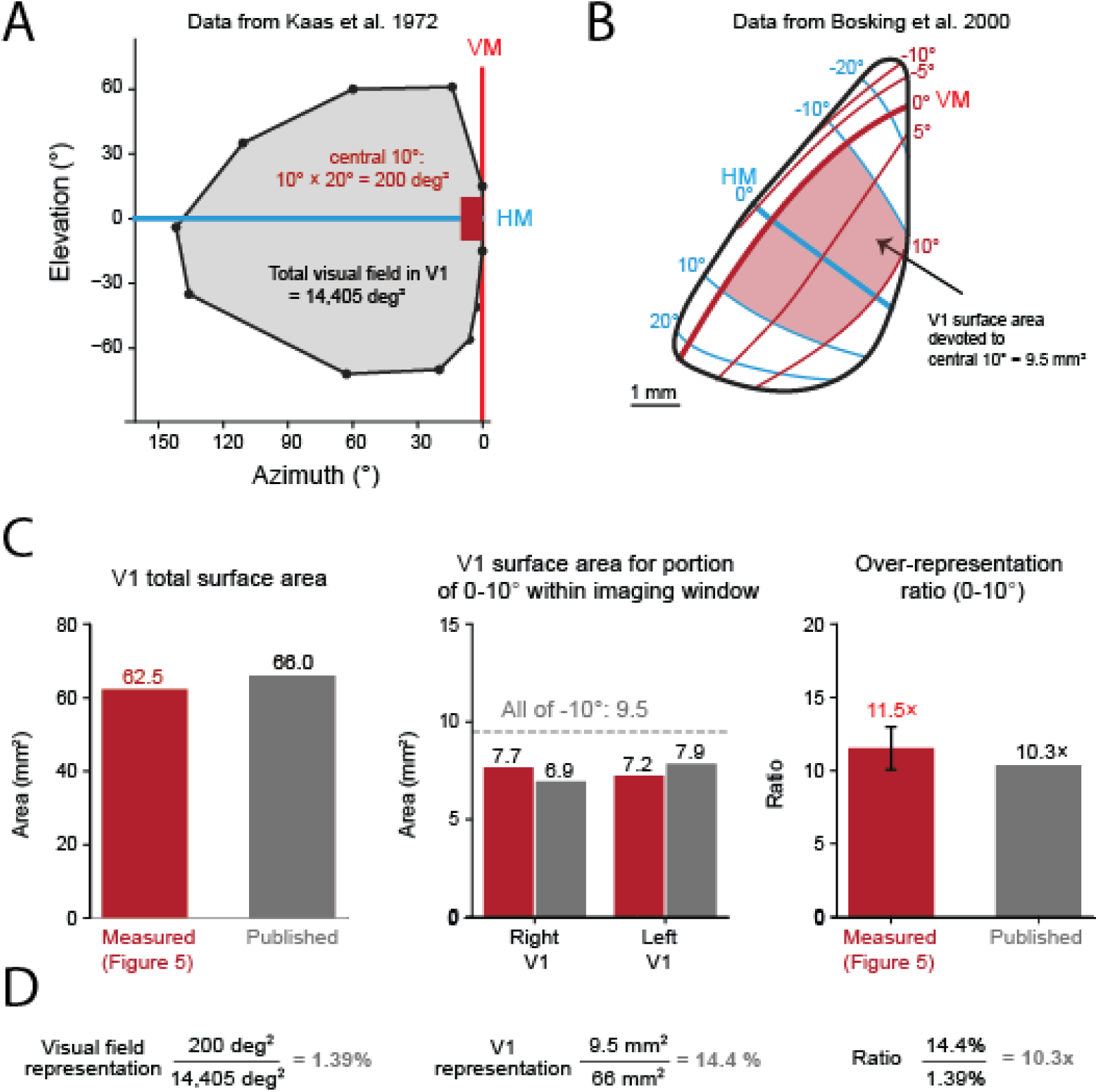
Supporting data for Figure 5. **Tree shrew over-representation ratio confirmed from published data. A)** Contralateral visual field coverage in V1, generated from electrophysiological data in Kaas et al ^26^. The total visual field, 14,405 deg², is the area of this polygon. VM is vertical meridian and HM horizontal meridian. Red rectangle depicts the central 10 degrees, which runs from 0-10 degrees azimuth and -10 to 10 degrees elevation, for a total of 200 deg². **B)** Retinotopic map of the dorsal part of V1 (left hemisphere), modified from Bosking et al.^25^. The red region represents the central 10 degrees, mapping onto the red region in A. The surface area of this region is calculated using the scalebar, and comes to 9.5 mm^2^. **C)** Comparison of our measurements (red) with published estimates (gray). Left: Total V1 surface area from our reconstructed cortical flatmap compared with published values^28^. Middle: Cortical area representing the portion of the central 10° captured by our imaging window in each hemisphere, compared with estimates from the Bosking et al.^25^ map. For this comparison, the published retinotopic grid was interpolated across a triangular mesh, and cortical areas were summed only for regions representing the same visual-field locations sampled by our imaging data. Right: Over-representation ratio calculated from our widefield measurements compared with an independent estimate derived from published cortical and retinotopic maps. **D)** We step through the over-representation ratio calculation for the published data.

**Figure S6:**
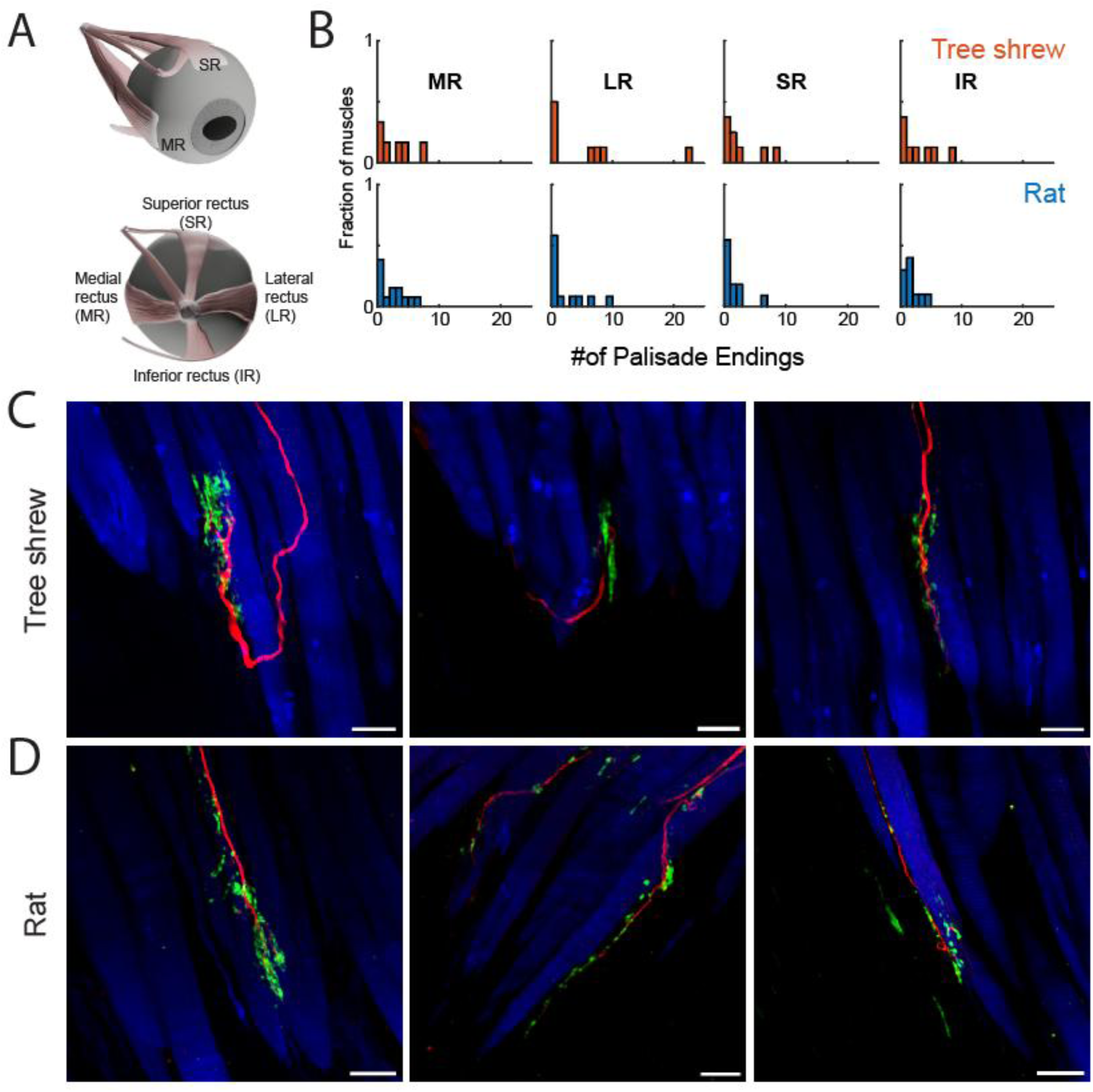
Supporting data for Figure 6. **A)** Schematic of the eye showing the four rectus muscles: superior (SR), inferior (IR), medial (MR) and lateral (LR) rectus. SR/IR contribute to vertical gaze shifts whereas MR/LR contribute to horizontal gaze shifts. **B)** Distribution of palisade ending counts per muscle for each rectus muscle in tree shrews (top, orange; n = 6 MR, 8 LR, 8 SR, and 8 IR muscles from 4 animals) and rats (bottom, blue; n = 13 MR, 12 LR, 11 SR, and 10 IR muscles from 7 animals), shown as the fraction of muscles. **C)** Additional example high magnification confocal z-projected images of palisade endings in tree shrew EOMs, labeled for neurofilament (NF, red), synaptophysin (SYP, green) and phalloidin (blue). Synaptophysin-positive terminals cluster at the end of the axon, around the muscle fiber tip. **D)** Same as C for rat EOMs. Synaptophysin-positive terminals are more sparse in rats and also distributed along the axon rather than clustered at the end as in tree shrews. Scale bars, 25 µm.

